# Partial sex linkage and linkage disequilibrium on the guppy sex chromosome

**DOI:** 10.1101/2022.01.14.476360

**Authors:** Suo Qiu, Lengxob Yong, Alastair Wilson, Darren P. Croft, Chay Graham, Deborah Charlesworth

## Abstract

The guppy Y chromosome has been considered a model system for the evolution of suppressed recombination between sex chromosomes, and it has been proposed that complete sex-linkage has evolved across about 3 Mb surrounding this fish’s sex-determining locus, followed by recombination suppression across a further 7 Mb of the 23 Mb XY pair, forming younger “evolutionary strata”. Sequences of the guppy genome show that Y is very similar to the X chromosome, making it important to understand which parts of the Y are completely non-recombining, and whether there is indeed a large completely non-recombining region. Here, we describe new evidence that supports a different interpretation of the data that suggested the presence of such a region. We analysed PoolSeq data in samples from multiple natural populations from Trinidad. This yields evidence for linkage disequilibrium (LD) between sequence variants and the sex-determining locus. Downstream populations have higher diversity than upstream ones (which display the expected signs of bottlenecks). The associations we observe conform to predictions for a genome region with infrequent recombination that carries one or more sexually antagonistic polymorphisms. They also suggest the region in which the sex-determining locus must be located. However, no consistently male-specific variants were found, supporting the suggestion that any completely sex-linked region may be very small.

## Introduction

Genetic sex determination and sexual dimorphism were first studied in the guppy, *Poecilia reticulata* (formerly *Lebistes reticulatus*) just over 100 years ago (Schmidt, 1920), and the species has been considered a good system for understanding the lack of recombination between sex chromosome pairs. This fish has male heterogamety, though like the sex chromosomes of many other fish, the XY pair is homomorphic, or, at most, slightly heteromorphic, with the Y being larger than the X (Lisachov, Zadesenets, Rubtsov, & Borodin, 2015; Nanda et al., 2014; Traut & Winking, 2001). Recent genome sequence analyses have shown that the gene contents of this chromosome pair are very similar, unlike the situation in many species (which often show extreme differences between the Y chromosome sequences and their X-linked counterparts). It has nevertheless been proposed that recombination suppression events have occurred repeatedly, producing extensive non-recombining regions, or “evolutionary strata” (Wright et al., 2017) like those in heteromorphic sex chromosome pairs, including those of humans and other mammals (Cortez et al., 2014; Lahn & Page, 1999), birds (Xu et al., 2019) and the plant *Silene latifolia* (Papadopulos, Chester, Ridout, & Filatov, 2015).

It is important to understand whether the guppy XY pair indeed has evolutionary strata, for the following reason. Trinidadian guppy Y chromosomes carry sexually antagonistic (SA) male coloration factors that are advantageous in males, and increase male mating success, outweighing the disadvantage of being more conspicuous to predatory fish that are present along with guppies in their natural habitats (Endler, 1980; Houde, 1992). These factors are polymorphic within Trinidadian guppy populations (Haskins, Haskins, McLaughlin, & Hewitt, 1961). Because the presence of SA polymorphisms is the most plausible situation that can generate selection for closer linkage with the sex-determining locus, resulting in evolution of suppressed recombination (Rice, 1987, see also a recent review by Otto, 2019), the guppy is ideal for studying this central hypothesis for the evolution of potentially extensive Y-linked regions.

The proposed guppy strata include an old stratum between 22 and 25 Mb in the assembly of chromosome 12 (LG12), which includes the sex-determining locus, which is thought to be located roughly 30% of the physical distance from the terminal region of the roughly 26.5 Mb telocentric XY pair (Lisachov et al., 2015; Nanda et al., 2014; Tripathi, Hoffmann, Weigel, & Dreyer, 2009). The younger strata are claimed to cover about 7 Mb of the more centromere-proximal part of the XY pair (Bergero, Gardner, Bader, Yong, & Charlesworth, 2019; Charlesworth, Bergero, Graham, Gardner, & Yong, 2020; Darolti, Wright, & Mank, 2020; Fraser et al., 2020; Wright et al., 2017). However, the Y has also been shown to cross over occasionally with the X chromosome, so that most of the chromosome is partially, rather than completely, sex-linked (reviewed by Lindholm & Breden, 2002), although recombination rates are very low across most of the chromosome. Based on molecular markers, crossovers have only rarely been detected in a large region from the centromere end to about 25.5 Mb (Bergero et al., 2019). Here, we describe new evidence that supports an alternative interpretation of the results that suggested that the guppy has recently evolved new evolutionary strata.

Although studies of guppy samples consistently yield evidence that guppy Y-linked sequences differ from their X-linked counterparts, the signals are weak, and strata containing many genes are not clearly defined by analysis including F_ST_ between the sexes, or divergence between Y and X sequences (Bergero et al., 2019; Charlesworth et al., 2020; Darolti et al., 2020; Fraser et al., 2020; Wright et al., 2017). It is likely that the male-determining region is physically small (Bergero et al., 2019), and it remains uncertain whether there is any extended completely sex-linked region. No completely male-specific variant (indicating complete Y-linkage) has yet been found, though all the studies just cited describe hints that associations between variants and the sex-determining locus are commonest in the region distal to 20 Mb of the chromosome.

In testing for such associations and attempting to understand neutral molecular variation in natural populations, such as those of guppies, it is important to take account of the demographic history of the populations sampled. Specifically, bottlenecks cause loss of low frequency variants, reducing diversity, and also increase the variance in diversity values. Bottlenecks can therefore lead to regions of high F_ST_, even when no selective process has affected the sequences. In samples of the two sexes, regions of the sex chromosome pair distant from the sex-determining locus may show high F_ST_ values, and other genome regions may appear as outliers with high values. Linkage disequilibrium (LD) between sequence variants and F_ST_ between two subdivided populations are equivalent (Charlesworth, Nordborg, & Charlesworth, 1997). In the context of subdivision into X- and Y-linked chromosome regions, such associations can be used to identify the location of the sex-determining locus. Sequences very closely linked to such polymorphisms maintained by long-term balancing selection should display associated neutral variants (in an XY system, male-specific variants). The region will therefore have higher sequence diversity in males than females. This signal might be particularly clear in bottlenecked populations, with low diversity elsewhere in the genome. However, stronger associations are expected in such populations, as small sample sizes result in strong LD by chance, even between sequence variants that are not closely linked (Park, 2019). Large samples will therefore be needed to detect the signal of Y-linkage in the face of randomly generated LD, and to distinguish between such LD and complete sex linkage (evolutionary strata). Given that bottlenecks probably occurred in the formation of populations of Trinidadian guppies in upstream locations (Magurran, 2005), their effects must be considered in drawing conclusions based on neutral molecular variation, particularly for upstream samples, which were inferred to have evolved evolutionary strata not found in downstream samples (Bergero et al., 2019; Charlesworth et al., 2020; Darolti et al., 2020; Fraser et al., 2020; Wright et al., 2017).

The effect on diversity of neutral variants linked to a balanced polymorphism in a population of constant size depends on the quantity r = 4*N*_e_*r*, where *r* is the recombination rate in males between a neutral site and the selected locus. Under commonly observed recombination rates, the effect is therefore restricted to a physically small region, but to larger regions if crossovers are rare, or *N*_e_ is very small. Deterministic results for the coalescence time between Y- and X-linked sequences when the selected locus is a sex-determining locus (Kirkpatrick, Guerrero, & Scarpino, 2010) are illustrated in Figure 1. Pairs of X-linked sequences should also show a peak of differentiation at the sex-determining locus, though this is much smaller than between Y- and X-linked sequences (Figure 1). Variants across larger physical distances can show associations with individuals’ sexes if a polymorphic SA factor is present in a partially sex-linked region closely linked to the sex-determining locus (Kirkpatrick & Guerrero, 2014). This effect might be detectable in the guppy if most of the Y recombines rarely with the sex-determining locus.

**Figure 1.**
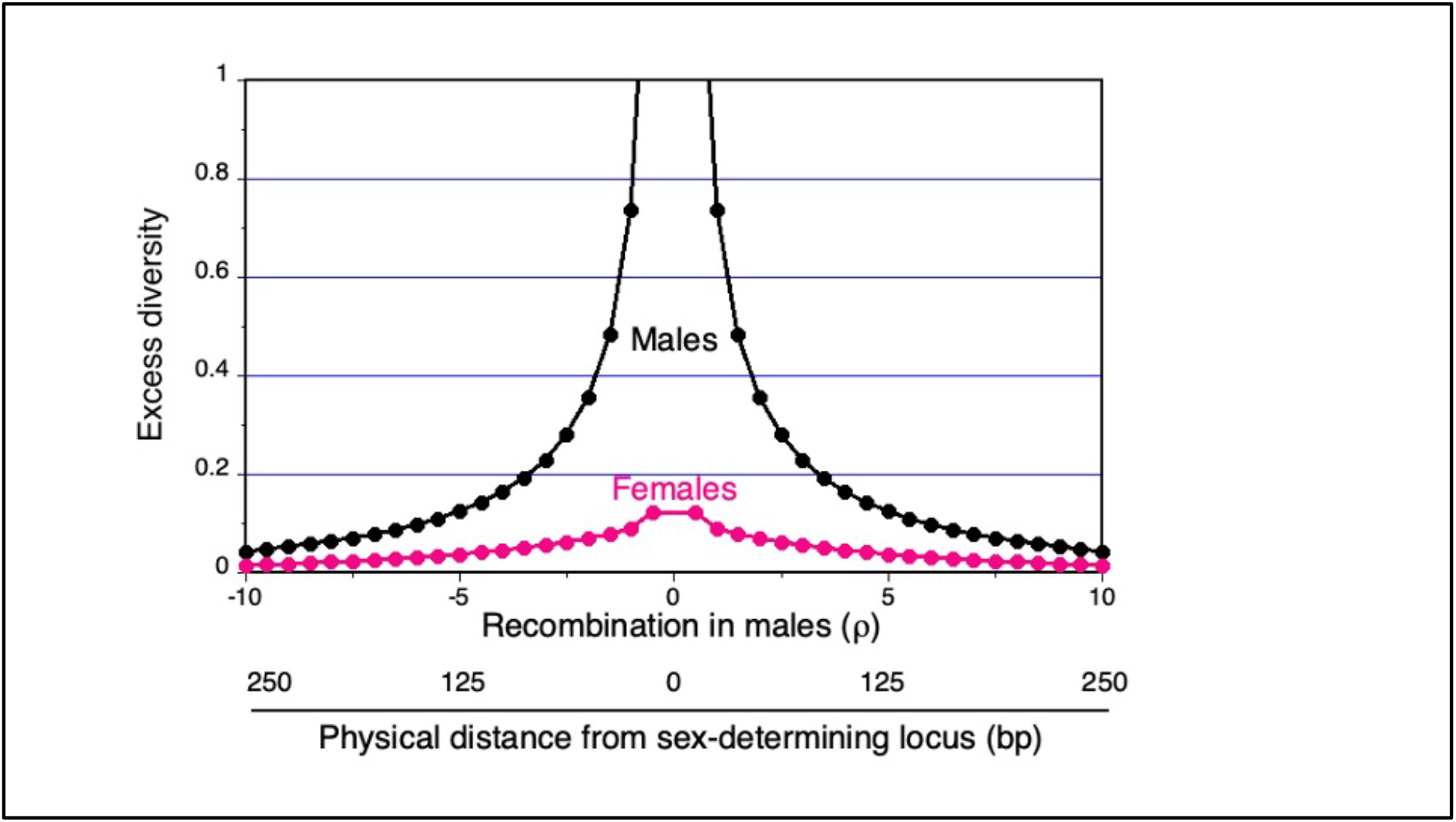
Peak of neutral diversity predicted in males and females at regions linked to a male-determining factor for the case with no sexually antagonistic polymorphisms in the region. The values are based on equations for coalescence times between pairs of X-linked sequences and between Y- and X-linked ones (Kirkpatrick, Guerrero, & Scarpino, 2010). The elevated Y-X divergence times can be detected in phased data as higher Y-X divergence values at neutral sites close to the male-determining factor, or in unphased data as higher diversity in males than females. The y axis shows the excess in diversity at different distances from a male-determining factor, compared with unlinked sites. Distances are shown on the x axis as the recombination rate in males, ⍰, or 4Nr values, where r is the recombination distance in Morgans), and also as the physical distances assuming a very low recombination rate of 0.01 cM/Mb and an N value of 1 million. A higher recombination rate would proportionately decrease the physical distances. Coalescence times (and consequently diversity values) are also elevated for X chromosome sequences, but only slightly.

Populations of guppies with the SA polymorphisms already mentioned are present in most rivers in the Northern Range mountains of Trinidad. Some of the polymorphisms have probably been maintained for long times, as they are shared by many different populations. They are found in both upstream sites, under low predation pressure, and downstream ones, with heavy predation (reviewed in Magurran, 2005), and in Tobago (Haskins et al., 1961). Their maintenance involves rare male advantages (BA Fraser, Hughes, Tosh, & Rodd, 2013; Hughes, Houde, Price, & Rodd, 2013; Olendorf et al., 2006), not solely SA selection. Nevertheless, the maintenance of SA polymorphisms leads to selection for the evolution of suppressed recombination, although this is not certain to occur, as the male coloration factors are also male-limited in expression (reviewed in Haskins et al., 1961). Differentiation between the sexes for sequences on LG12, which carries the guppy sex-determining locus, was inferred to be stronger in samples from upstream than downstream sites, suggesting evolution of suppressed recombination (Wright et al., 2017).

Although the effects of different predation regimes on the maintenance of partially sex-linked SA coloration factors, and the evolution of suppressed recombination, have not been modelled, the ecological differences between up- and downstream sites might allow sex chromosome strata to evolve only in upstream sites. One possibility is that, in downstream sites, recombinant females that inherit and express coloration factors will be selectively disadvantaged, keeping the frequency of these factors so low that selection against recombinants might be too weak to select for closer linkage with the male-determining factor.

An alternative is that suppressed recombination between the XY pair should evolve most readily in downstream sites, because of the high predation rate that would affect recombinant females that inherit and express coloration factors. This view is supported by evidence for more frequent (though still rare) recombination between a sex-linked male coloration factor (*Sb*) and the male-determining locus in up-than downstream males from the Aripo river (Haskins et al., 1961). Moreover, stronger associations between the male-determining locus and the *Sb* factor in high-than low-predation males, using testosterone treatment to reveal the factors present in females, were supported by consistent results in other rivers for coloration (Gordon, López-Sepulcre, & Reznick, 2012). These results suggest that low recombination rates between the coloration factors and the male-determining locus prevail in high-predation conditions, but, after colonisation of upstream sites, with weaker predation pressure, recombinants are less disfavoured (reviewed in Charlesworth, 2018). This hypothesis implies that close linkage in high-predation populations is reversible, and not caused by a Y chromosome inversion. Indeed, the genome sequences of the male and female of this species from a high-predation site in the Guanapo river show no rearrangement between the Y and the X (Fraser et al., 2020; Künstner et al., 2017). Instead, the recombination rate difference between the two kinds of populations seems more likely to be due to a shift in the localisation of crossovers, or in the crossover rate in an LG12 region that recombines rarely in males from high-predation sites, consistent with changes detected in evolutionary experiments in the guppy (Gordon, López-Sepulcre, Rumbo, & Reznick, 2017).

The conclusion that evolutionary strata have evolved only in low-predation populations is further called into question by evidence that crossovers occasionally occur in the regions identified as younger strata (Almeida et al., 2021; Bergero et al., 2019; Charlesworth et al., 2020; Darolti et al., 2020). Even rare crossing over will prevent accumulation of Y-specific variants, and indeed, as mentioned above, no consistently male-specific variants have so far been found in guppy populations. It is therefore worth considering alternative explanations, such as the bottleneck hypothesis outlined above, for the finding that regions showing associations with the sexes extend across larger regions in samples from low-predation populations, compared with other populations.

Here, we compare diversity and F_ST_ values between the sexes for the XY pair and the autosomes for Trinidadian guppy population samples from several rivers. Although autosomal diversity was reported by Almeida et al. (2021) only relative X/A and Y/A values were interpreted. However, it is important to consider the absolute values, because high F_ST_ values can be due to low diversity, since F_ST_ reflects the proportion of total diversity that is found between, rather than within populations, and low diversity within one or both of the populations being compared necessarily implies high F_ST_ (Charlesworth, 1998; Charlesworth et al., 1997; Cruickshank & Hahn, 2014). This can also affect F_ST_ values between the sexes. For example, if an advantageous X-linked mutation has recently spread through a population, causing a selective sweep, the resulting low diversity of X chromosome alleles could create high male-female F_ST_.

Our aims were as follows:

1. To analyse genome-wide patterns of synonymous site diversity to help understand natural Trinidadian guppy populations and test for recent bottlenecks in upstream populations.
2. To test whether the region in which the guppy male-determining factor is known to be located displays unusually high nucleotide diversity, relative to values in other genome regions in individuals sampled from the same population, or whether the entire sex chromosome shows elevated diversity, or whether parts of the X chromosome might show evidence for low diversity (due to possible selective sweeps on the X), which could account for regions of high male-female F_ST_ values.
3. If the entire sex chromosome shows elevated diversity in males, to test whether females also show higher diversity on this chromosome pair, compared with the autosomes, as predicted above if the elevated diversity in males is due to LD with the sex-determining locus under partial sex linkage.
4. To examine the sex chromosome region in which SNP genotypes are most strongly associated with individuals’ phenotypic sexes, to narrow down the location of the guppy male-determining factor, and to assess the likely size of any non-recombining region, or evolutionary stratum. In the Discussion section, we discuss problems that may be responsible for the difficulty experienced in locating this factor.

## Methods

### Trinidadian guppy samples

Fish were sampled from 12 natural populations in Trinidad (Supplementary Table S1), and 20 males and 20 females were preserved in the field before transport to the UK for DNA extraction and sequencing.

### Mapping pooled sequences to Guppy female assembly

Low coverage Illumina sequencing was done for separate male and female pools, each of 20 individuals, for each sample, yielding sequence lengths of 150 bp, and the raw sequence reads from the 24 pools were processed as described previously (Yong, Croft, Troscianko, Ramnarine, & Wilson, 2021). Briefly, reads were trimmed to remove low quality bases and adapter sequences using *cutadapt*. The cleaned reads were mapped to the guppy female assembly v1.0 (accession number: GCF_000633615.1) using BWA (H. Li & Durbin, 2010). Mapped reads were sorted and PCR duplicates were removed using Samtools v1.9 (H Li et al., 2009). Reads around indels were realigned using the modules RealignerTargetCreator and IndelRealigner implemented in gatk v3.4. Finally, the input file for the diversity analyses described in the next section was generated using the Samtools mpileup function, reads with mapping quality lower than 20 were discarded, and SNPs were called in Popoolation.

### Nucleotide diversity analyses in males and females

To ensure that the sequences used are reliably aligned single-copy genes, and the variants analysed are likely to behave close to neutrally, we used synonymous sites in annotated genes for population genomic analyses of nucleotide diversity. Synonymous nucleotide diversity (π_S_) and Watterson’s ϴ_S_ were estimated using the Popoolation software (Kofler et al., 2011); the software’s Syn-nonsyn-sliding.pl and Syn-nonsyn-at-position.pl scripts were used to estimate these quantities in non-overlapping 250Kbp windows and in individual genes, which yielded similar results (see below). To obtain reliable diversity estimates, SNPs with coverage lower than 1/3 of the average coverage for the sample, or higher than 3 times the average coverage, were first filtered out. SNPs for these calculations were defined with one as the minimum number of reads supporting the minor allele, to check that the π_S_ results are consistent with those using the SNPGenie software (Nelson, Moncla, & Hughes, 2015). As described later, the conclusions based on diversity values were not changed by setting this number to 2 (mc=2 in Popoolation; reads with alternative variants below that threshold are assumed to be errors). Because ϴ_S_ estimates depend on rare variants (the majority of variants in populations), the threshold needed in Popoolation analyses affects these values, as discussed in the Results section. Diversity values were estimated separately for the samples of males and females from each collection site. Finally, values of the quantity 1 - (π_S_/ϴ_S_), denoted by Δϴ_S_, were calculated for each window or gene. This detects frequency differences between variants, as values of ϴ are estimated using variants irrespective of their frequencies, whereas π values largely reflect intermediate frequency variants. Δϴ_S_ can therefore detect the excess of rare variants that is expected in a population as it returns towards the neutral equilibrium frequency distribution after a recent bottleneck; in such populations, higher values of Δϴ_S_ are expected (Tajima, 1989).

The Δϴ_S_ measure is preferable to Tajima’s D values as it is less affected by the lengths of the sequences analysed.

### PoolSeq detection of candidate fully sex-linked sites and sex-linked regions

To test for candidate completely sex linked variants on the sex chromosome pair, we screened the data from each sample of 20 males and 20 females to detect sites with variants consistently showing the genotype configuration expected under complete sex linkage: under male heterogamety, the guppy sex-chromosome system (Winge, 1922b), all females will be homozygous for an X-linked variant at candidate sites, while all males will be heterozygous for a different variant. Male-specificity in a large enough sample indicates Y linkage, though not necessarily complete Y linkage (see discussion below and Supplementary Figure 1). Sites with the configuration suggesting XX in females and XY in males are called “XY sites” in what follows. This analysis used all site types, not just synonymous sites, because variants of any kind can help detect sex linkage. Previous analyses using Illumina sequencing data (Almeida et al., 2021; Bergero et al., 2019; Charlesworth et al., 2020; Reichwald et al., 2009) have also analysed regional differences in the proportions of polymorphic sites with genotype configurations suggesting different strengths of association with sex-determining loci, but sample sizes were small, or the individuals were from captive populations. Large samples from multiple natural populations are needed to narrow down the location of the guppy sex-determining locus, and our study adds new samples to those analysed previously (Almeida et al., 2021; Bergero et al., 2019; Charlesworth et al., 2020), and describes the results of the analyses in greater detail, revealing valuable new information (see Results).

Table 1 illustrates our approach for such screening of PoolSeq data, based on the frequencies of alternative bases at biallelic sites. At a site where a variant was detected in a sample of females, the proportion of the variant (XFREQ) was computed using the total number of reads of either base, after the filtering described above (TOTAL). This detects candidate sites with no variation in the X chromosomes present in the females. At all such sites with variants detected in the male sample, we calculated the proportion of the variant, multiplied by 2, yielding YFREQ, the variant’s estimated frequency in the population of Y chromosomes. The table illustrates the method with three sites. When all 20 males are heterozygotes for a base that differs from the one found in all females, the YFREQ value is 1, suggesting fixation of the variant in the Y population (as expected for the male-determining locus, which must be present in all males). YFREQ values of 1, suggesting fixation in the Y population, may also be found at sites completely linked to the male-determining factor.

**Table 1.** Examples of the method for finding candidate fully sex-linked variants among variants detected in the sample from a natural population of a species known to have a genetic sex-determining system with male heterogamety. The commonest variant in the females (in bold font in the table), is defined as the X-linked allele and the alternative variant at the same site is inferred to be Y-linked. The total coverage at a site is denoted by “TOTAL”.

| Site |  | Counts of A:T:C:G at a site |  | XFREQ=<br>X/TOTAL | YFREQ=<br>Y/TOTAL*2 |
| --- | --- | --- | --- | --- | --- |
| Position in the<br>female assembly | REF<br>base | Females | Males |  |  |
| 12 | T | 0:**33**:0:0 | 0:**29**:0:1 | 1 | 0.017 |
| 9320 | T | 2:**44**:0:0 | 23:**48**:0:0 | 0.957 | 0.648 |
| 9414 | G | 14:0:0:**88** | 105:0:0:**75** | 0.863 | 1.167 |

Because the individuals were sequenced in pools of males and females, the true numbers will be less than 20 of each sex. It is nevertheless unlikely that many variants will be at high intermediate frequencies in the males, and have this YFREQ value, but yet be invariant in the female sample. For example, consider a site with both XFREQ and YFREQ values of 1, which thus has an estimated frequency of the rarer variant (detected only in the males) of 0.25 (see Table 1). Most sites satisfying this criterion will be XY sites that are likely to be sex linked, even if many fewer than 20 females were in fact sequenced. For instance, if half of the individuals of either sex carry the allele detected in the males, the binomial probability that the variant is not present in a female pool of only 10 individuals is 0.098% (ignoring genotyping errors in the male sample, or failure to detect the variant in heterozygous females). A further problem is that sequencing or mapping errors (or rare recombination events) might produce XFREQ values slightly below 1 in some samples, which would not be included as candidates for having male-specific variants. We therefore repeated the analysis with XFREQ ≥ 0.975 (i.e. one of the 40 alleles in our female sample matching the putatively Y-specific allele). As described below, this did not increase the sharing of candidate fully sex-linked sites between populations.

This approach has the potential to detect candidate sites with patterns suggesting complete sex linkage, and chromosome regions that show enrichment for such sites. Importantly, some variants at sites within a completely sex-linked region may have YFREQ values < 1, because a recent mutation in a Y-linked sequence may not have become fixed at the site (although, in the absence of recombination, such a segregating variant will still be male-specific, and the site will have XFREQ = 1). Segregating male-specific variants might also be due to maintenance of different Y haplotypes. We consider this in the Discussion section, along with comparing this approach with alternative methods used for screening for sex-linkage. Variants in partially sex-linked regions may also have high YFREQ values, but both these, and their XFREQ values should be < 1.

## Results

### Nucleotide diversity in different populations and evidence for recent population size bottlenecks

With both diversity measures we estimated (see Methods), most low-predation (LP) populations have the expected lower diversity than the high-predation (HP) ones from the same river (Figures 2 and 3 and Table 2), for both LG12 and autosomal sequences (except for the Quare river samples, individually labelled in Figure 3, with no diversity difference); the patterns are similar for diversity based on genes, rather than windows (Supplementary Table S3A). Even including the Quare river, mean π in upstream populations is at least 37% lower than in downstream ones, based on autosomal sequences from both sexes (or 41% based on medians); this diversity measure is conservative for detecting diversity loss, as π depends most strongly on variants at intermediate frequencies, which are least likely to be lost. The differences in θ are larger (Supplementary Table S3A), as this diversity measure is more weighted to low frequency variants (see Methods), which are most likely to be lost in bottleneck events.

**Figure 2.**
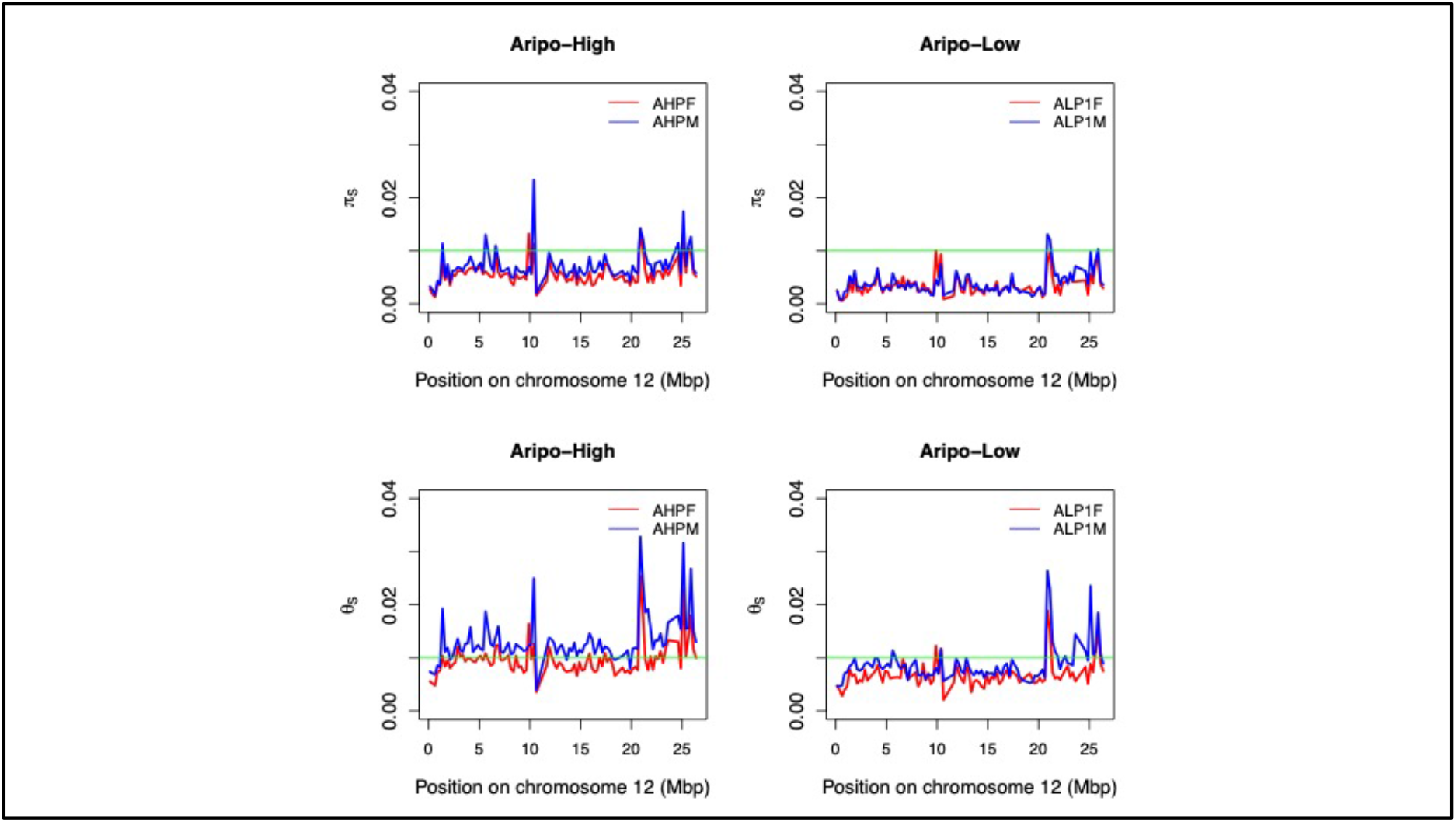
Example of diversity estimates from the Aripo river, showing the two diversity measures, both based on synonymous sites (plots for all samples analysed are in Supplementary Figure S2). The values shown are mean values in 250 kbp windows, with red for female samples and blue for male ones. The green lines indicate a diversity level of 1%, to emphasise the lower diversity in in the low-than a high-predation collection site (left- and right-hand plots, respectively), and its much higher θ_s_ than π_s_ value, suggesting a recent bottleneck in this site.

**Figure 3.**
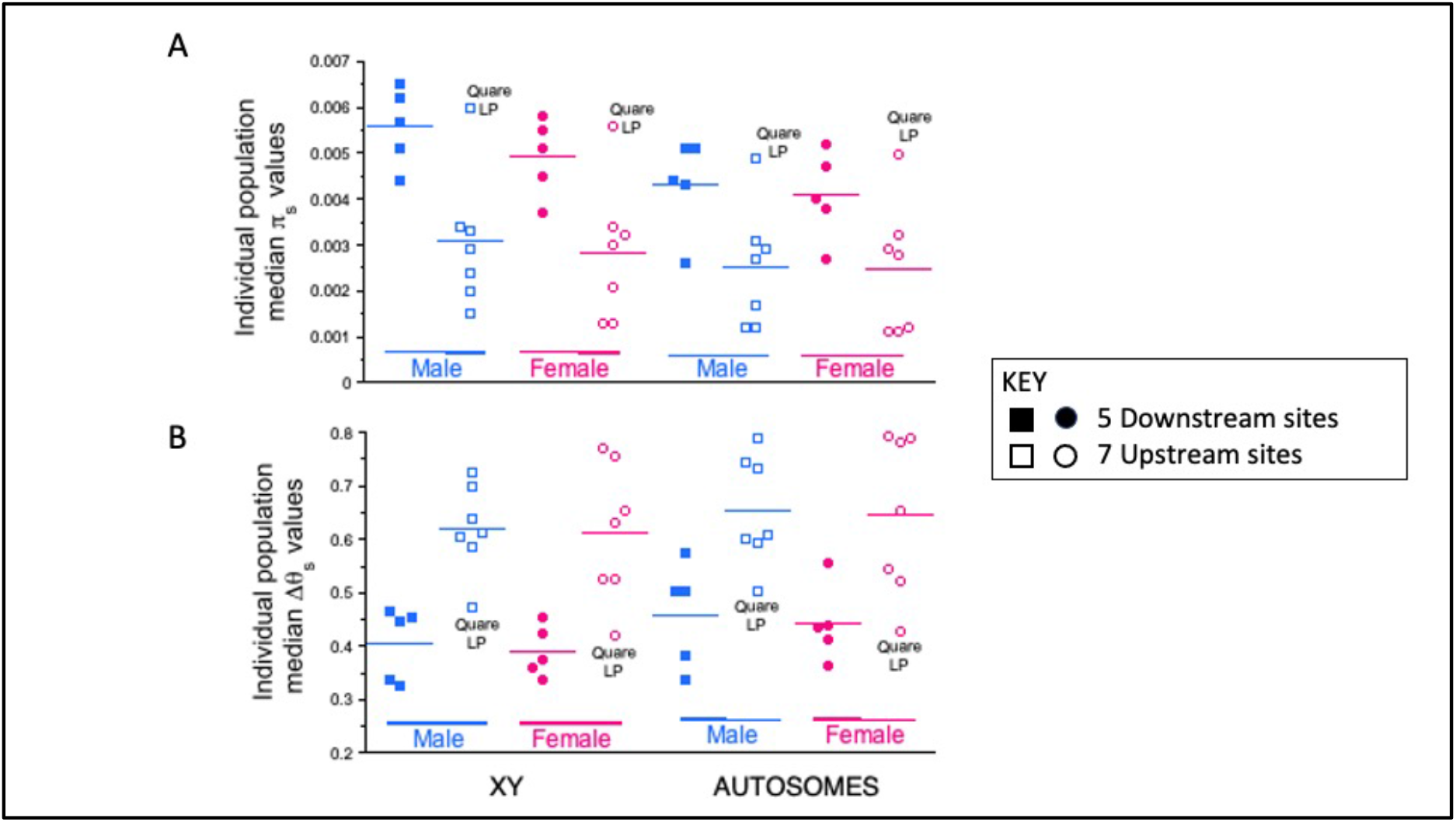
Diversity estimates in all populations sampled. Part A shows the consistently lower values of synonymous site diversity (π_s_) in low-predation upstream sites (solid symbols), compared with downstream, high-predation collection sites from the same rivers (open symbols), and part B shows the higher Δθ_s_ values in the former, suggesting that upstream sites have consistently been affected by recent bottlenecks. Blue and pink symbols show male and female results, respectively.

Both high- and low-predation samples also show deficits of low-frequency variants (Δθ_s_ values, see Methods, are shown in Figure 3 and Supplementary Table S3). Because the sequences were obtained by PoolSeq, absolute values of this quantity cannot be interpreted, but the values are consistently higher in three of the four rivers where up- and down-stream samples could be compared, with significant differences in both sexes, for both autosomal and sex-linked variants (Supplementary Table S4). Three rivers have only one predation regime represented (Guanapo, Paria and Petit Marianne, see Supplementary Table S1), and the results from these are also consistent with this pattern (Figure 3, Supplementary Tables S3 and S4). However, the Quare upstream sample (Quare LP in Figure 3) is again unusual, as its unexpectedly high diversity (for both LG12 and autosomal sequences) is accompanied by an absence of evidence for loss of rare variants, and its Δθ_s_ value is low, like the values for downriver samples, indicating an abundance of rare variants. The differences between up and downstream samples were largely unchanged with a higher threshold number of reads supporting the minor allele (see Methods), though this reduced the diversity estimates; as expected, since many sites with low frequency minor alleles were filtered out, θ_s_ was greatly reduced, but upstream samples still consistently had high Δθ_s_ values, compared with downstream ones (Supplementary Figure 3B and Table S3C).

The lower diversity of the upstream samples is expected to lead to comparisons involving such samples having the highest F_ST_ values. This can explain the high values in comparisons between pairs of upstream populations from different rivers, using all site types, and lower ones between upstream and downstream populations from the same river (B. A. Fraser, Künstner, Reznick, Dreyer, & Weigel, 2015; Suk & Neff, 2009; Willing et al., 2010) (the same is seen for the samples and synonymous sites studied here, using the PoolFstat package (Hivert, Leblois, Petit, Gautier, & Vitalis, 2018), Supplementary Table S2.

### Male-female differentiation

Figures 2 and Figure 3A show that we also detect higher synonymous site diversity in males than females. The sex difference is especially clear in downstream samples where diversity his high (Figure 3 and Supplementary Figure S3; Supplementary Figure S2 shows detailed results for LG12 for all 12 populations sampled). Again, these differences were maintained under a higher threshold number of reads supporting the minor allele (Supplementary Table S3C). Because separate pools were sequenced from males and females, and sex differences in diversity were not confined to the sex chromosome (Supplementary Table S3B), we quantified the effect of sex linkage using the ratio of sex chromosome/autosomal diversity values in each sex. Almost all samples have excess diversity on the sex chromosome (Figure 4, Supplementary Figures S2B and S3). Interestingly, this is detected in both sexes, not just in the males, though the excess is larger in males than females in 9 of the 12 samples. In all high-predation populations, and in the low-predation Quare sample, excess diversity was detected in males across much of chromosome 12; all these samples had at least 70 windows with ratios above 1 (out of the total of 106 250 kb chromosome 12 windows), highly significantly exceeding the expected 53 windows under the null hypothesis that this chromosome has diversity values similar to those for autosomal sequences. Among the female samples, only the high-predation one from the Guanapo river yielded a significant excess of such windows (69/106). This population also showed high F_ST_ between males and females in a previous study (Fraser et al., 2020).

**Figure 4.**
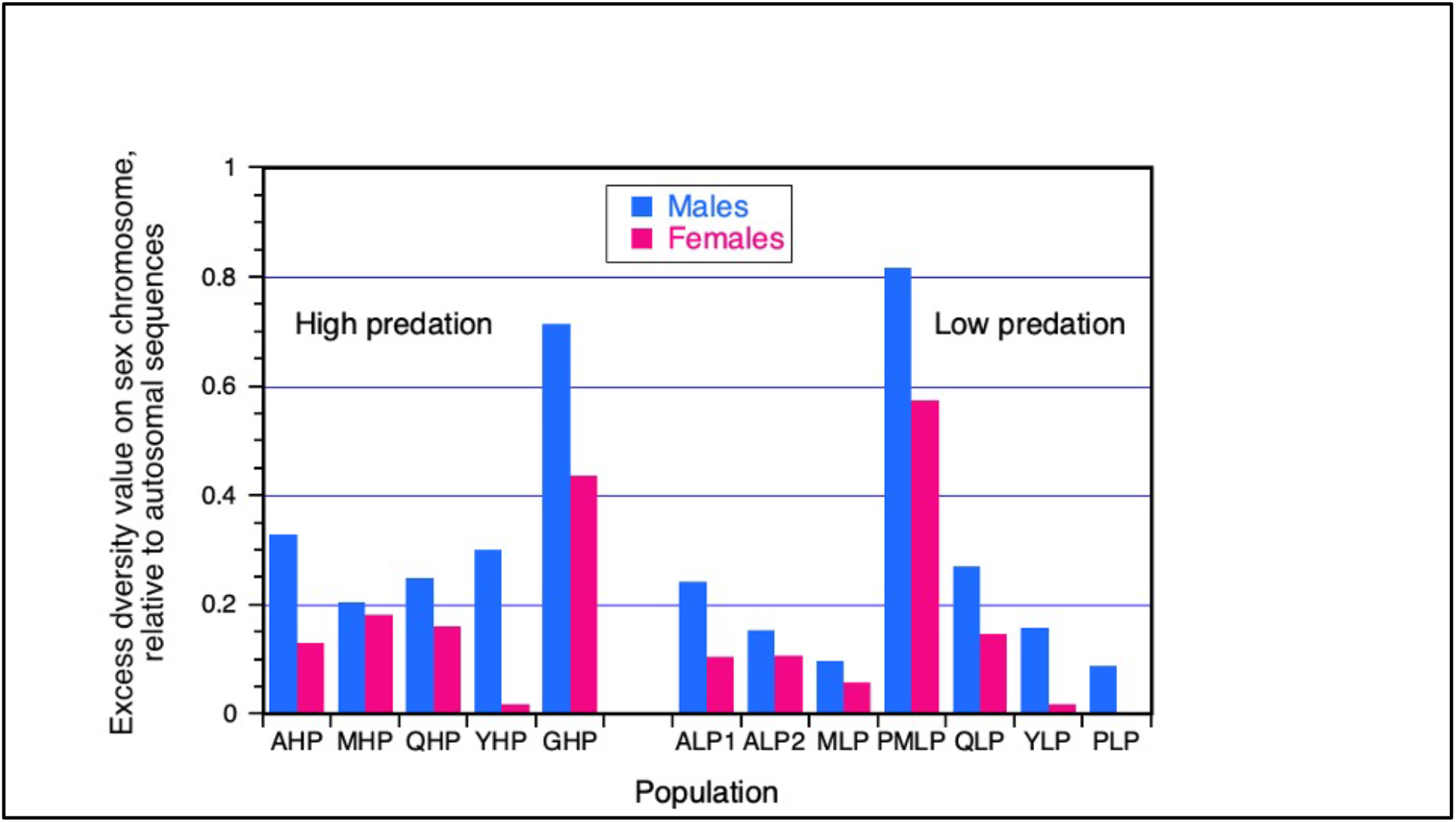
Excess of the estimated diversity on LG12, the sex chromosome, versus the autosomal estimates. The nucleotide diversity estimates are for synonymous sites (values of πS), based on results analysed in windows (see Methods). For each population sample shown on the x axis, the plot shows values of the extent to which the LG12/autosome diversity ratio exceeds 1. Blue bars show the results for males and pink bars those for females.

The tendency for LG12 sequences sampled from females to have higher diversity than autosomal ones suggests that the X chromosome has not experienced recent hard selective sweeps. Genetic maps of LG12 in female guppies are consistently around 50 centiMorgans (Bergero et al., 2019; Charlesworth et al., 2020), implying that more than a single crossover event per meiosis is rare. A recent strong selective sweep event would therefore create a wide region of very low diversity (Supplementary Figure S2).

### Attempts to locate the guppy male-determining region

The LG12 region in which the guppy male-determining factor is known to be located (distal to 20 Mb) appears to show higher synonymous site diversity in males than females, unlike the rest of the chromosome (Figure 2, Supplementary Figure S2). However, the difference is small, and the coverage is low in the region, and, across LG12, other high diversity values are clearly associated with low coverage (Supplementary Figure S8). Overall, these appears to be repetitive regions, and may not reliably identify the region that includes the male-determining factor. We therefore used a different approach, based on associations between sex and genotypes at variable sites on LG12. The approach searches the PoolSeq data for sites (of any type, not just coding region sites) where the genotypes in males and females conform to those expected under complete sex linkage, as described in the Methods section, which also describes the criteria used for including a site in the searches. Figure 5 shows results for sites that were polymorphic in each population, and at which variants were detected only in the population’s male sample (while the females had good coverage, but were monomorphic, with XFREQ values equalling 1), and had YFREQ values above 0.9, indicating strong association with the male-determining locus (see Methods). Although male-specific variants are concentrated in the expected region, the picture for the entire LG12 (Figure 5) does not point clearly to a completely sex-linked region; the upper part of Figure 5 shows that at most 4 populations have sites satisfying this stringent Y criterion. This is consistent with the diversity results described above, and with the previously published results described in the Introduction. Supplementary Figures S4 and S5A show results for male-specific variants with YFREQ = 0.7, to allow for non-fixation in the Y chromosome population. This does not alter the regions showing most associations with the sex-determining locus. If we include sites where one allele in our female sample matched the putatively Y-specific allele (see Methods) the sharing of candidate fully sex-linked sites between populations did not increase (Supplementary Figure S5B).

**Figure 5.**
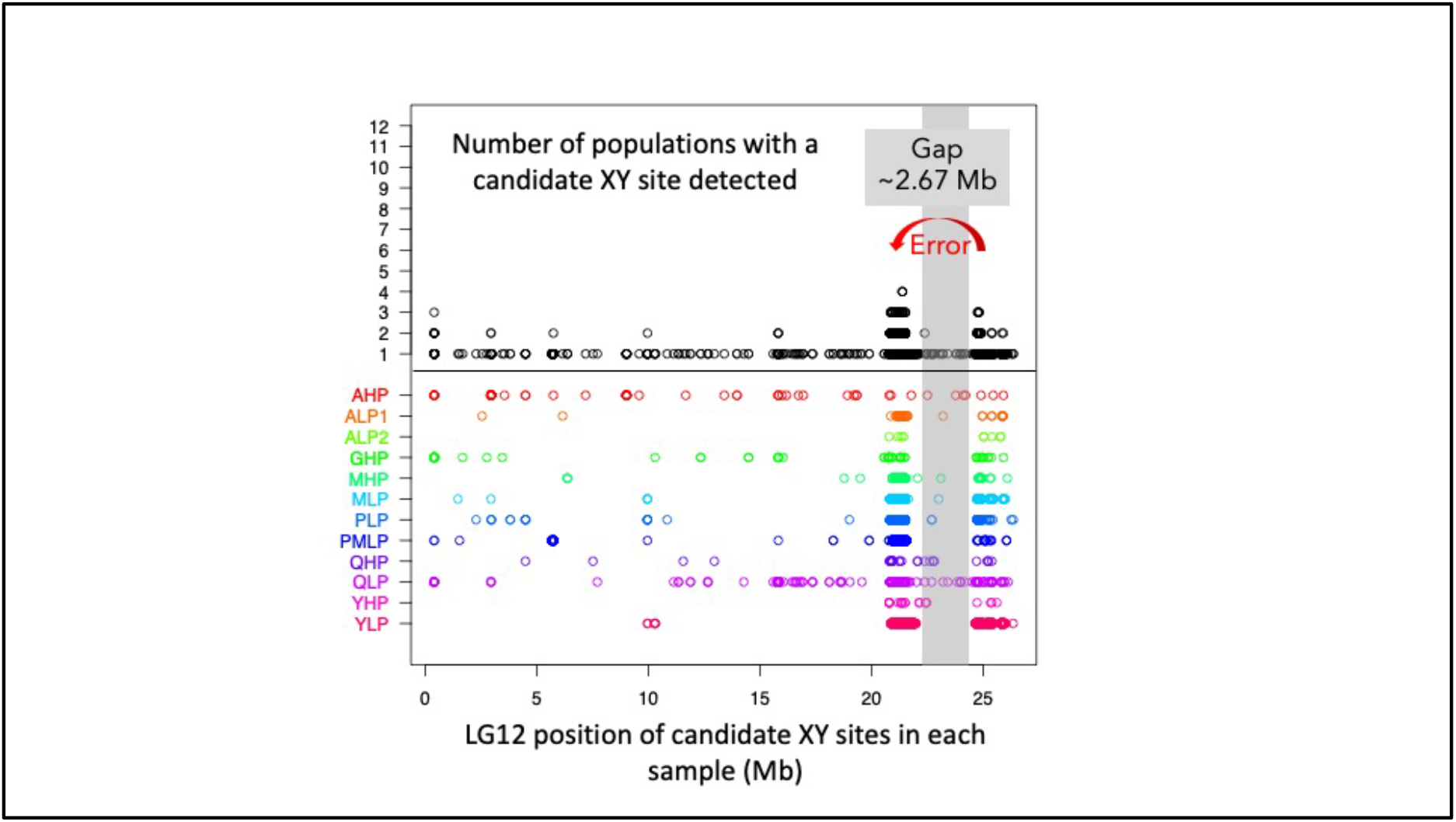
Candidate fully sex-linked sites in the entire LG12, based on PoolSeq data. SNPs at all site types (synonymous or non-synonymous) were analysed in our samples of 20 individuals of each sex from each of 12 natural populations, and the plot shows results for sites with no variants in the 20 diploid sequences from females (XFREQ = 1 in Table 1). The upper section shows the results for each such site that varied within any of the populations analysed, and the y axis shows the number of populations where a site satisfied the criterion to be classed as having the XY configuration, suggesting that it is a good candidate for complete sex-linkage. The analyses results shown used the criterion that the YFREQ value was at least 0.9, indicating that at most two males in the sample of 20 could have lacked the putatively Y-linked alternative variant at the site. No site has a male-specific candidate SNP in all 12 populations. The bottom part shows the locations of candidate male-specific SNPs in each of the individual population samples (in different colours), to show the consistent signals in the terminal part of the chromosome in all the different populations.

Overall, these analyses do not definitively identify either the male-determining locus or any clear fully sex-linked region in which it is located. Of the total of 7,705 candidate sites with male-specific variants in at least one population sampled, 79% were detected in only one of our 12 samples. The broad signals in some samples may reflect the expected wide LD in bottlenecked populations explained above. However, some high-predation samples (including those from the Aripo and Guanapo rivers) yielded few candidate sites, and very few were detected in the Aripo LP2 sample (though their locations are consistent with the region identified in the other samples, Supplementary Figure S4). Instead of a clear sex-linked region, two regions of the sex chromosome, between roughly 20.5 and 21.8 Mb, and distal to 24.5 Mb in the female assembly, are enriched in sites with apparently male-specific variants, while the signal is much weaker between 21,519,986 and 25,369,837, with no candidates shared by more than 2 populations. No recombinants have yet been found between markers within these three regions and the male-determining locus (Charlesworth et al., 2020), and they may all be completely sex-linked, or almost so, while the pseudo-autosomal region or PAR starts more distally (slightly centromere-proximal to 25,194,513 bp, see (Charlesworth et al., 2020); the regions are shown, together with information about their genetic map locations, in Supplementary Figures S6, and Supplementary Figure S7 shows that the two sub-regions with the strongest signal of sex linkage also have high repetitive content. The gap between these two regions does not reflect low coverage (Supplementary Figure 8). The terminal parts of the sex chromosome may still not be correctly assembled, given their high repetitive content and presence of sequences that were unplaced in the male assembly (Supplementary Figures S7 and S6), and the correct location of gap region could be distal to the two regions with male-specific SNPs.

Only 36 of the potentially male-specific sites satisfy the threshold of YFREQ ≥ 0.7 in at least 4 population samples, 29 of which are distal to 20.9 Mb. The candidate site found in the largest number of populations (at position 21,410,653) is found in only 7 samples. Excluding two sites distal to the gap region suggests a candidate sex-linked region with a total size of 590,830 bp, which includes only 22 genes (Supplementary Table S5). Two, cyclin and shroom3-like, are in the previously proposed candidate region (contig IV gene island), whose repetitive parts had higher coverage in males than females, and are duplicated near 24 Mb in the assembly of the sequenced male (Fraser et al., 2020). Another candidate region (Dor et al., 2019), in the more terminal LG12 region assembled distal to the gap, was also detected in this study of fish from natural populations, but no candidate genes emerged (Fraser et al., 2020).

## Discussion

Our analyses indicate a greater genome-wide excess of rare synonymous site polymorphisms in upstream than downstream guppy populations, indicating that at least the upstream ones are not at mutation-drift equilibrium. This will pose problems for population genomic approaches aimed at detecting footprints of balancing selection (as here) or adaptive changes.

Low diversity in upstream populations is consistent with previous studies of polymorphic microsatellite or SNP markers (Barson et al., 2009; Willing et al., 2010).and very high F_ST_ values between pairs of samples from upstream populations from different rivers, though previous studies of F_ST_ either did not provide nucleotide diversity estimates (Suk & Neff, 2009; Fraser et al., 2015) or did not detect bottlenecks (Willing et al., 2010). F_ST_ estimates using the same PoolSeq data as analysed here (Yong et al., 2021 see Supplementary Table S2), support the same pattern, and a recent population genomics study using all site types inferred the occurrence of bottlenecks in several rivers (Whiting et al., 2021). Our analyses of synonymous, probably weakly selected, variants in coding sequences reveals specific loss of rare variants, as expected if upstream populations are recovering after having lost diversity in their recent history. Their low diversity is therefore unlikely to reflect long-term low effective population sizes. It also cannot reflect extremely severe recent bottlenecks causing almost complete loss of variability, and leaving only variants at intermediate frequencies, which would cause a difference in Δϴ_S_ values opposite to that observed. Metapopulation dynamics, with local extinction and recolonisation events involving bottlenecks, could explain the observed low diversity in up-stream guppy populations (Figure 3A, Supplementary Table S3) and very high F_ST_ values between up-river sites from different drainages.

Rare migration from downstream populations seems unlikely to have produced the observed consistent pattern. Unless migration rates are very low, samples of more than 10 individuals per deme in a subdivided population should not display strong departures from equilibrium (Table 7.1 of Charlesworth & Charlesworth, 2010). Migration from upto downriver localities is probably more frequent (Whiting et al., 2021). The Quare results differ from the general pattern, possibly because of migration of fish from headwaters of other rivers during floods.

### Diversity within the two sexes, and diversity differences between sex chromosomes and autosomes

Synonymous sites, under weak selective constraints reducing diversity, are also preferable for testing for the predicted higher diversity in males than females from natural populations. We indeed consistently detect sex chromosome-specific differences (Figures 3 and 4), whereas a previous analysis of F_ST_ between guppy males and females, uaing all site types, did not detect clear differences between LG12 and the autosomes (Fraser et al., 2020).

Diversity is higher than for autosomal sequences across the entire guppy LG12 in downstream populations (Figure 2 and Supplementary Figure 2). Upstream populations might have been expected to show the clearest signals, because associations between SNPs and the male-determining locus (under balancing selection) should be maintained even if the rest of the genome has lost variability. However, LD generated by sampling during bottlenecks in these populations may outweigh this. Also, as outlined in the Introduction, the signal may be restricted to a small region near the male-determining locus, as in two fish whose sex-determining factors are allelic differences of single genes within recombining genome regions, the fugu (Kamiya et al., 2012) and species in the very distantly related genus *Seriola* (Koyama et al., 2019), and maybe also in *Nothobranchius furzeri*, another fish with homomorphic X and Y chromosomes (Reichwald et al., 2015).

The large LG12-wide excess of diversity in male guppies from high-predation populations suggests that the whole chromosome (other than the terminal pseudo-autosomal region) is generally transmitted from fathers to sons, with only rare crossing over. Although family studies cannot reliably estimate very low recombination rates, they show that rates in male guppies are very low across most of this chromosome (Bergero et al., 2019; Charlesworth et al., 2020; Haskins et al., 1961). Associations might thus be found across much greater physical distances than in fugu or *Seriola*, and male coloration factors might therefore be maintained at any of the many genes located within the rarely recombining region. A very low population recombination rate (r < 5) would be required to account for the observed almost 20% higher sex chromosome than autosomal diversity in males in the absence of a SA polymorphism (Figure 1). Presence of a SA polymorphism can, however, increase the Y-X coalescence times and at sites closely linked to the polymorphic locus, and also increases coalescence times between X-linked regions (Kirkpatrick & Guerrero, 2014). We indeed detect elevated X-linked, compared with autosomal, diversity in samples of females (Figure 4).

*P. wingei* appears to show greater LD, across a larger proportion of LG12, than *P. reticulata*. (Almeida et al., 2021; Darolti et al., 2020; Darolti et al., 2019), and this too could reflect associations with variants across a wide LG12 region. Samples from natural populations are not available, and the small sample sizes studied from this species will be even more important than in the guppy. Alternatively, the Y of this species may carry more male coloration factors.

### Approaches for discovering completely sex-linked regions and variants

Many approaches can detect an extensive non-recombining sex-determining region, or evolutionary stratum containing many sites with male-specific variants, will be detected by, but it will be difficult to detect a small region, or a non-recombining region that evolved recently, or one in which occasional recombination events still occur. Physically small male-determining loci, however, will be difficult to locate in genomes if associations arise due to close, but incomplete, linkage across large regions, especially if linked SA polymorphisms are maintained. Low diversity populations might then show the clearest difference between the sex-linked region and the rest of the genome, making the male-determining locus more detectable. In small or bottlenecked populations, LD is expected to be higher than larger ones, because lower r leads to greater LD at equilibrium (Haddrill, Thornton, Charlesworth, & Andolfatto, 2005). The footprint of sex-linkage might therefore be detected at more distant variants in such populations. On the other hand, bottlenecks increase the variance of LD, and result in strong LD by chance, even between sequence variants that are not closely linked (Park, 2019).

The presence of male-determining factor on the guppy LG12 could nevertheless potentially be detected by testing for regions with higher diversity, whether the functional site is in a coding or a non-coding region, using analyses of individual sites, as proposed here, to find variable sites with XY genotype configurations. Given a male-determining factor shared by all populations of a species, even a single base change from the sequence in females should be detectable. In the guppy, associations are consistently found within a region of the sex chromosome distal to 20 Mb, and our analysis (Figure 5) confirms that most of the associated SNPs lie within two regions, while a region between these largely lacks such SNPs (Almeida et al., 2021). Our analysis detected somewhat clearer associations in downstream than upstream samples (Figures 4 and 5).

The approach used here may be preferable to using F_ST_ values between the sexes. F_ST_ can detect sites associated with the sex-determining locus (Almeida et al., 2021; Bergero et al., 2019; Fraser et al., 2020). However, very high F_ST_ values can arise if diversity in one of the populations is low, since F_ST_ quantifies the proportion of diversity found between populations (Charlesworth, 1998; Charlesworth et al., 1997; Cruickshank & Hahn, 2014). When the sexes are treated as two populations, this could happen if there has been a recent selective sweep on the X chromosome, causing low diversity in females, or a recent bottleneck that eliminated much X-linked diversity leaving a high proportion of the remaining diversity due to Y-X divergence, even if that divergence is small in absolute terms. Such events could explain the chromosome-wide high F_ST_ values between males and females observed in a Guanapo population sample, but not other populations (Fraser et al., 2020).

Furthermore, if distinct Y haplotypes co-exist, each with “private”, haplotype-specific variants (as might occur if the rarely recombining region carries polymorphic male coloration factors), this will lead to low F_ST_ between the sexes. Diverse Y haplotypes have been uncovered in *Poecilia* (=*Micropoecilia*) *parae* (Sandkam et al., 2021), a species with polymorphic Y-linked male coloration (Lindholm, Brooks, & Breden, 2004). Three different male colour patterns are associated with distinctive k-mers, indicating that some sequence variants are present only in males with certain colour phenotypes, and supporting complete Y linkage (Sandkam et al., 2021). In such a situation, the male-determining factor should be shared by all males, but variants in the completely linked region will have arisen independently on the haplotype of each coloration factor, and many sites could show variants that are not shared with other males (Supplementary Figure S1). Analysing individual sites, rather than F_ST_ in windows containing multiple variable sites, partially avoids this problem, because only shared Y-linked variants that arose before the different Y haplotypes split will be detected. However, if individuals are sequenced separately and can be phased reliably, LD analysis can potentially resolve such situations more completely.

Our approach can nevertheless fail to find completely sex-linked variants, or physically small sex-linked regions. If the reference genome assembly of a species with male heterogamety is derived from sequencing a female, and if the male-determining gene is a Y-specific sequence, such as a duplication with a male-determining effect, it will not be represented in the reference genome. This is unlikely in the guppy, as use of a male genome assembly also failed to reveal clear associations (Fraser et al., 2020). Second, regions of high repeat density, may prevent Y- and X-linked sequences aligning reliably, resulting in gaps even if the male-determiner is a single nucleotide polymorphism in a gene present in both the X- and Y-linked regions. Our filtering to removed regions with high coverage (see Methods), may have created gaps where repetitive sequences are abundant, potentially excluding sites with XY genotype configurations. However, the absence of strong associations between about 21.8 and 24.5 Mb appears not simply to reflect this problem, as coverage values are not very low in this region (unlike parts of the flanking regions with high densities of male-specific SNPs, Supplementary Figure S8).

The two chromosome 12 regions could reflect separate peaks of variation associated with two loci under balancing selection: the male-determining factor, and a sexually antagonistic polymorphism, or cluster of polymorphisms, as modelled by Kirkpatrick & Guerrero (2014).

Another possibility that cannot currently be excluded, is that the different populations have different male-determining factors (Almeida et al., 2021). Non-1:1 primary sex ratios should then arise in crosses between or within populations. Sex ratio data are scarce in guppies. To our knowledge, data are not yet available from newly born progeny or unborn fry extracted from gravid females (which could be obtained using microsatellite or other molecular markers located in the sex chromosome region that recombines rarely, and therefore behave as Y-linked in most progeny of heterozygous males). However, inter-strain crosses and crosses involving YY males showed that female-biased sex ratios in laboratory strains of guppies are determined primarily by Y-linked genes decreasing production or competitive ability of Y-bearing sperm, possibly reflecting accumulation of deleterious alleles, or pleiotropic effects of Y-linked alleles that increase male fitness (Farr, 1981).

Finally, perhaps the genome region is arranged differently in different individuals. This could explain the different location of the signal in the Quare samples (though the signal at 20.9 Mb is weak).

Male-specific variants proximal to 20 Mb probably mainly reflect LD due to infrequent recombination. Some parts of the guppy sex chromosome pair recombine very rarely with the male-determining factor, while others do so at rates of up to 8%, though it is unclear whether these rates occur in natural populations, as many genetic studies used captive ornamental fish (Lindholm & Breden, 2002). It is unclear whether exchanges of coloration factors between the Y and X are observed because these factors are located in the region of frequent recombination distal to the male-determining locus, or reflect crossovers proximal to this locus. It is also unknown whether crossovers occur nearer the centromere end of the chromosome, where cytogenetic studies of crossover locations (Lisachov et al., 2015) and genetic mapping of molecular markers (Charlesworth et al., 2020) suggest that they are very rare.

## Conclusions

Overall, our results show that there is no need to invoke complete sex linkage, or the evolution of new non-recombining strata, to explain the pattern of differentiation between males and females across different parts of the guppy LG12. There may be no extensive (multi-gene) completely non-recombining region corresponding to evolutionary strata of other organisms such as humans. The appearance of strata may reflect the LD that is expected across regions that rarely recombine with the male-determining locus, and especially in bottlenecked populations. The male-determining locus is probably within a region 600 kb near 21Mb in the guppy female LG12 assembly, much smaller than the 3 Mb suggested oldest stratum (Wright et al., 2017). This is consistent with evidence that the male-determining factor maps distal to a marker at 21.3 Mb, while the boundary of the pseudo-autosomal region is near 25 Mb (Charlesworth et al., 2020). Although the assembly of the terminal part of LG12 remains uncertain, the sex-linked region could be confined to a single SNP in or near one of the 22 genes in the region defined here (Supplementary Table S5).

The localization of crossovers at one end of each chromosome arm in male guppies, may be an ancestral state, as emerging results from sex-specific genetic maps suggest similar patterns in other fish (Sardell & Kirkpatrick, 2019). Stronger terminal localization of recombination in male guppies may have evolved, but there is currently no evidence for such a change. Therefore, rather than selecting for suppressed recombination between loci with SA polymorphisms and the male-determining locus, it is plausible that low recombination across most of LG12 (producing close linkage with the male-determining factor), favoured establishment of such polymorphisms. The resulting associations of variants across the XY pair (Kirkpatrick & Guerrero, 2014) would resemble young evolutionary strata.

LD across the sex chromosome could reflect presence of other SA polymorphisms not just male coloration factors. Guppy populations show several other sexual dimorphisms, including smaller size of males than females and behavioural differences (Olendorf et al., 2006). However, it is currently unclear whether the differentiation detected across the guppy sex chromosomes implies the presence of multiple SA factors, as theoretical modelling shows that this requires strong selection at loci that are tightly linked, but distant from the sex determining region, and the loci remain polymorphic only if selection favours heterozygotes (Otto, 2019). The evolutionary history that determines nucleotide diversity depends on recent demographic changes, not just on the recombination rates and effective population sizes (i.e. r values in Figure 1) under which populations are evolving. Upstream populations have undergone such changes, and also evolutionary changes since these sites were colonised, as reviewed by Magurran (2005), and selective sweeps may have affected diversity of some genome regions. Therefore, even if the low recombination rates across much of the guppy LG12 could be estimated, it may be impossible to use the fit of diversity results to theoretical equations to test whether LD with weakly selected synonymous variants is so high that SA polymorphisms must be invoked to account for it.

## Acknowledgements

This project was supported by ERC grant number 695225 (GUPPYSEX). We thank Mateusz Konczal (Adam Mickiewicz University, Poznań, Poland) for helpful suggestions about analyses.

## Data Accessibility and Benefit-Sharing

Genomic data (raw fastq files) are available under ENA Project Accession PRJEB45804. “Benefits Generated: Collaborating scientists from the University of the West Indies, Trinidad, R. Mahabir and R. Heathcote assisted with the collection of the natural population samples and Trinidad-UK shipment of guppies, respectively, as acknowledged in Yong. et al (2021).

Benefits Generated: Benefits from this research accrue from the sharing of our data and results on public databases as described above.”

## Author Contributions

designed research: DC

performed research: LY, AW, DPC

analyzed data: SQ, LY, CG

wrote the paper: DC, AQ, AW, DPC

## Supplementary files

### Supplementary Tables

**Table S1**. List of samples and locations in Trinidad.

**Table S2**. F_ST_ between samples from different populations using all site types.

**Table S3**. Diversity in the two sexes in the samples from different populations, estimated for synonymous sites, showing lower diversity in upstream low-predations sites than in downstream sites with high predation, except in the Quare river. Note that the Guanapo river is represented by only a high-predation sample, the PM one only by a LP site, and the Paria river has no high-predation sites.

**Table S4**. Significance values of Wilcoxon tests for differences in the values of the quantity Δθ_s_ (see main text) between samples from high- and low-predation sites from 4 rivers where samples were available from both site types. The Δθ_s_ values were estimated using synonymous sites.

**Table S5**. Genes within the LG12 region with the most candidates for complete sex linkage.

### Supplementary Figures

**Figure S1**. Haplotypes in a genome region linked to a male-determining factor, such that genotypes of some sites in the region are associated with the sex phenotype. The columns on the left diagram three X haplotypes and 6 Y ones, and the rows show the states at 12 hypothetical variable sites, with blue colours indicating mutations that arose since the male-determining factor arose in the region (sites that have retained the same state in the Y and X haplotypes are not shown). At least one variant, the male-determining factor itself (dark blue), is by definition, found only in males (and shows complete association with maleness, assuming complete penetrance and no environmental effects. This factor could be an SNP in a gene, or a duplication into the region. Pink indicates the X-linked allele at this site and other sites.

Part A shows the 5 configurations possible when there is a single Y haplotype. Subsequent mutations in a completely Y-linked region will be male-specific (barring repeated mutation at the same site in the Y and the X), but will initially be rare among the Y haplotypes, and show partial associations with the sex phenotype phenotype (configuration number 3); they may later become fixed fixed in the population of Y haplotypes, and thus become completely associated with the sex phenotype (configuration number 2). In the diagram, only one site other than the sex-determining locus is completely associated with this factor, with all Ys carrying the blue variant and all Xs carrying the pink one (configuration2; in the notation used in Table 1, such variants will have a frequency of 0.5 is males, and YFREQ value of 1, and a frequency of 0 in females, with XFREQ also = 1). Sites outside the completely sex-linked region may recombine onto the X, and show partial associations with the sex phenotype. Sites with configurations 4 and 5 must be partially sex-linked (and some with configuration 3 may also be in this category). Configuration 5 sites have a polymorphism shared by both sexes, and configuration indicates 4 sites that are fixed for a variant in the Y sample, but segregating in the X sample.

Part B shows the case when there are two Y haplotypes, so that variants in the completely sex-linked region are often not fixed in the sample of Y chromosomes. If the existence of two haplotypes was not recognised, these sites would be assigned configuration 3. If these are the majority of the variable sites showing associations with the the sex phenotype, it might be very difficult to determine the position of the maleness factor, especially if information is missing for some sites. Configuration 2 (if validated in a large enough sample) suggest sites in an ancestral Y haplotype that evolved before the separation into types 1 and 2.

**Figure S2**. Nucleotide diversity estimates (π_S_ values) for LG12 for all sampled populations (Figure 2 shows the example of Aripo river downstream and upstream samples), and the Δϴ_S_ values (see Methods). Part A shows the results without normalisation, and part B shows values of normalized by the values for autosomal genes estimated from the sample of the same sex.

**Figure S3**. Sex differences in estimated nucleotide diversity at synonymous sites (values of πS), based on results analysed in windows (see Methods), in samples from the sites whose names are shown above the plots. High-predation (downstream) sites’ names are in blue boxes, and those of low-predation, upstream sites are in green boxes. Results for females are shown in the two boxes outlined in red at the left of each plot and for males (blue outlines), on the right of each plot, and separate estimates were obtained for genes on the sex chromosome, LG12 (coloured pink boxes labelled S on the x axis) and on the autosomes (green boxes labelled A).

**Figure S4**. Associations between variants and individuals’ sexes for each population sampled, for the region distal to 21 Mb of chromosome 12. The results show sites with no variation in the female samples (XFREQ= 1, labelled XP=1 in the titles of each plot), and with a range of YFREQ values (labelled YP values in the key), from the most stringent value of 1, down to the value of 0.7 shown in Figure S5.

**Figure S5**. Candidate fully sex-linked sites in the entire LG12, analysed and displayed in the same way as in Figure 5 of the main text, but with YFREQ values > 0.7, instead of 0.9.

**Figure S6**. Comparison of positions of sites on the guppy sex chromosome in the male and female assemblies. The plot shows distal to the region that was inverted in the female assembly (see Figure S7), and it sites that have been genetically mapped to different regions in male meiosis are indicated in different colours. Markers within sequences in the gap region shown in Figures 5 and S5 are shown as open circles, and all markers mapped in this region show complete sex linkage in the sires of both families mapped. 25 markers map to the PAR in at least one of the sires studied, and 16 map to a region, contig IV, that is also duplicated at 5 Mb (Fraser et al., 2020). The plot also shows positions in the female assembly of several scaffolds that were unplaced in the male assembly, but that map genetically to this part of the guppy sex chromosomes in our sires.

**Figure S7**. Comparison of the female (x axis) and male (y axis) assemblies of the guppy sex chromosome. High densities of repetitive sequences can be seen at 10 Mb, and in both regions that include most of the male-specific SNPs (se Figures 5 and S5).

**Figure S8**. Mean coverage of sites in 10 kilobase windows across the sex chromosome in males and females in our pooled sequence data, with the windows where male-specific SNPs were detected shown in red (using the same threshold as for Figure S5, Yfreq=0.7), and other polymorphic sites in grey. The vertical dashed red lines indicate the two regions with high densities of SNPs showing associations with maleness; the coordinates for these regions (in the female assembly) are 20.5 - 21.8 Mb and 24.5 - 26.2 Mb. The figure shows the Marianne River low predation samples, as an example. Coverage results from other rivers are very similar.

