## Supplementary Figs for "Partial sex linkage and linkage disequilibrium on the guppy sex chromosome"

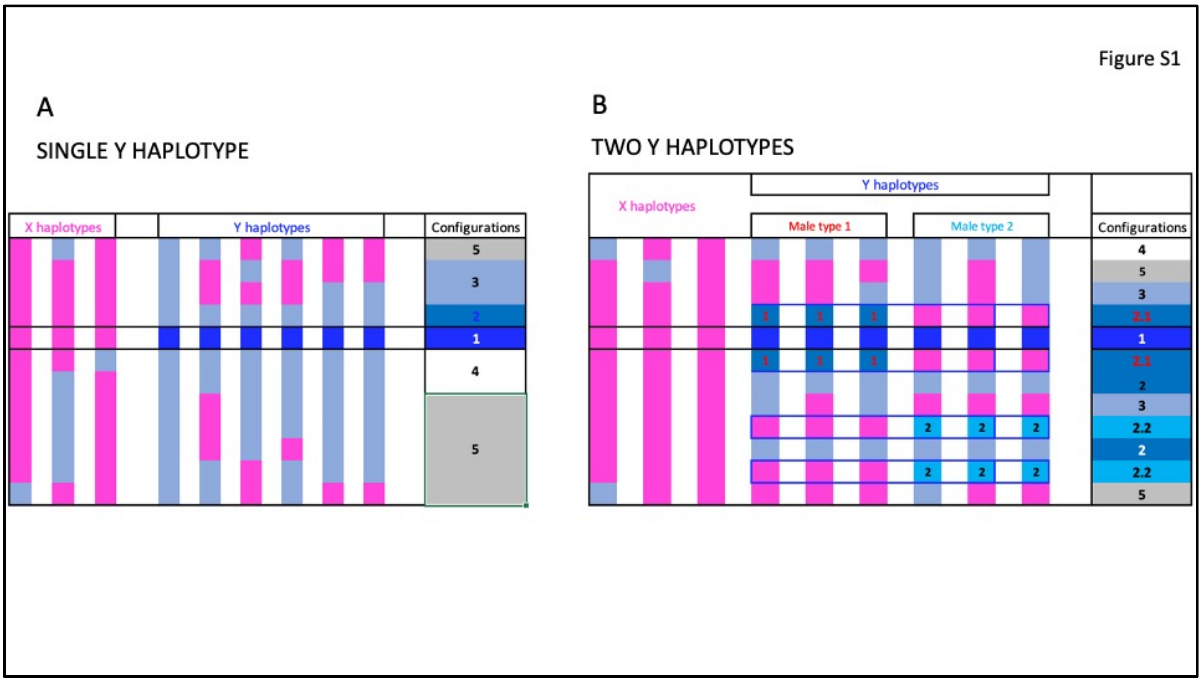


Figure S1. Haplotypes in a genome region linked to a male-determining factor, such that genotypes of some sites in the region are associated with the sex phenotype. The columns on the left diagram three X haplotypes and 6 Y ones, and the rows show the states at 12 hypothetical variable sites, with blue colours indicating mutations that arose since the male-determining factor arose in the region (sites that have retained the same state in the Y and X haplotypes are not shown). At least one variant, the male-determining factor itself (dark blue), is by definition, found only in males (and shows complete association with maleness, assuming complete penetrance and no environmental effects. This factor could be an SNP in a gene, or a duplication into the region. Pink indicates the X-linked allele at this site and other sites.

Part A shows the 5 configurations possible when there is a single Y haplotype. Subsequent mutations in a completely Y-linked region will be male-specific (barring repeated mutation at the same site in the Y and the X), but will initially be rare among the Y haplotypes, and show partial associations with the sex phenotype phenotype (configuration number 3); they may later become fixed fixed in the population of Y haplotypes, and thus become completely associated with the sex phenotype (configuration number 2). In the diagram, only one site other than the sexdetermining locus is completely associated with this factor, with all Ys carrying the blue variant and all Xs carrying the pink one (configuration2; in the notation used in Table 1, such variants will have a frequency of 0.5 is males, and YFREQ value of 1, and a frequency of 0 in females, with XFRE Qalso = 1). Sites outside the completely sex-linked region may recombine onto the X, and show partial associations with the sex phenotype. Sites with configurations 4 and 5 must be partially sex-linked (and some with configuration 3 may also be in this category). Configuration 5 sites have a polymorphism shared by both sexes, and configuration indicates 4 sites that are fixed for a variant in the Y sample, but segregating in the X sample.

Part B shows the case when there are two Y haplotypes, so that variants in the completely sex-linked region are often not fixed in the sample of Y chromosomes. If the existence of two haplotypes was not recognised, these sites would be assigned configuration 3. If these are the majority of the variable sites showing associations with the the sex phenotype, it might be very difficult to determine the position of the maleness factor, especially if information is missing for some sites. Configuration 2 (if validated in a large enough sample) suggest sites in an ancestral Y haplotype that evolved before the separation into types 1 and 2.

**Figure S2**. Nucleotide diversity estimates ($\pi_S$ values) for LG12 for all sampled populations (Figure 2 shows the example of Aripo river downstream and upstream samples), and the $\Delta\Theta_S$ values (see Methods). Part A shows the results without normalisation, and part B shows values of normalized by the values for autosomal genes estimated from the sample of the same sex.

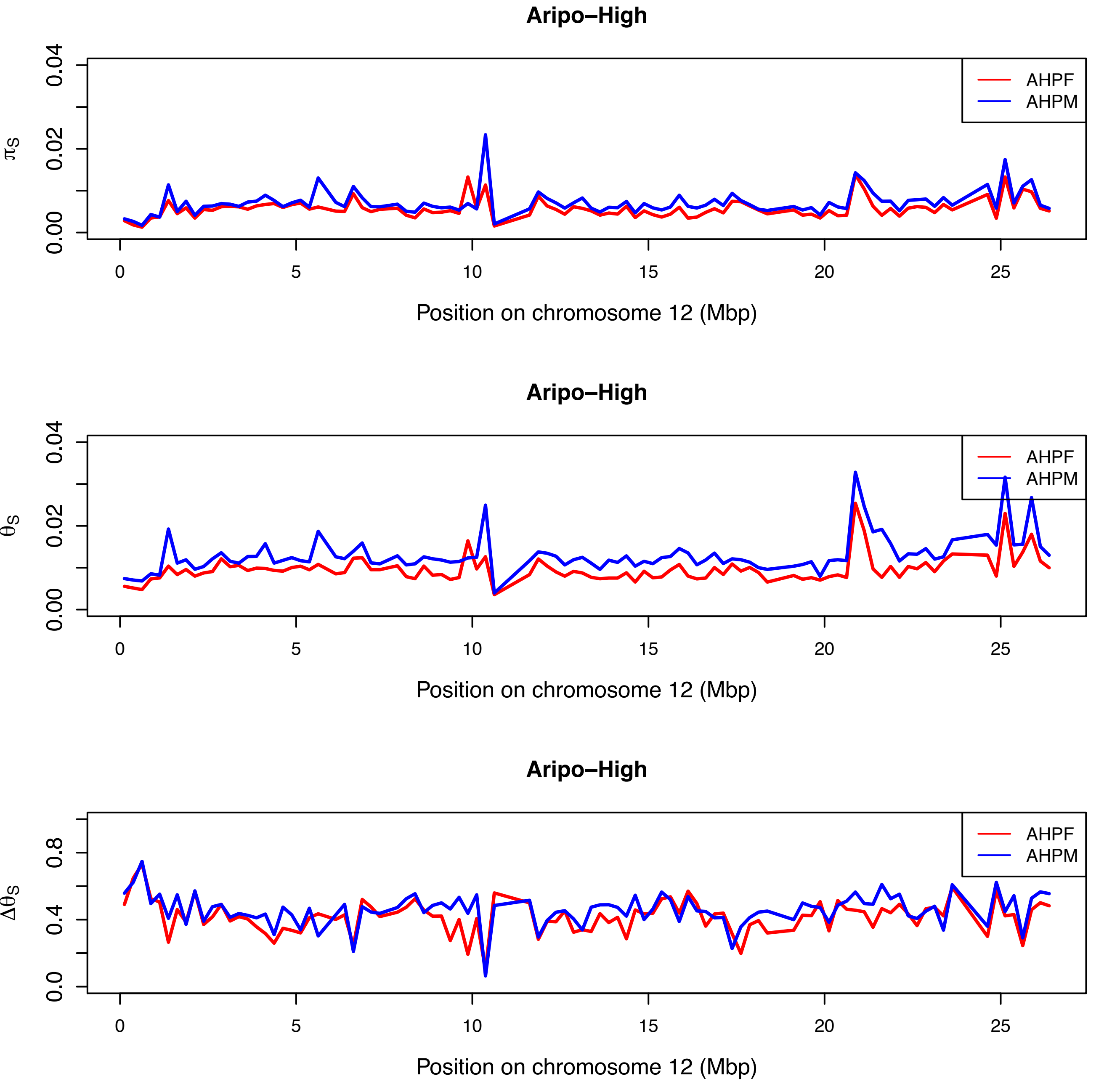
Aripo–High
AHPF
AHPM
πS
0.00
0.02
0.04
0
5
10
15
20
25
Position on chromosome 12 (Mbp)
Aripo–High
AHPF
AHPM
θS
0.00
0.02
0.04
0
5
10
15
20
25
Position on chromosome 12 (Mbp)
Aripo–High
AHPF
AHPM
ΔθS
0.0
0.4
0.8
0
5
10
15
20
25
Position on chromosome 12 (Mbp)

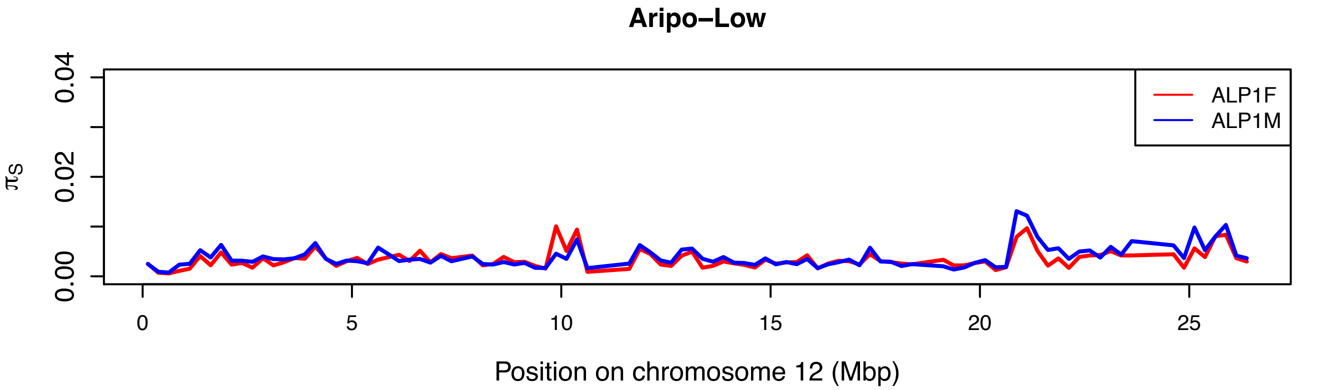
Aripo–Low
ALP1F
ALP1M
$\pi_S$
0.00
0.02
0.04
0
5
10
15
20
25
Position on chromosome 12 (Mbp)

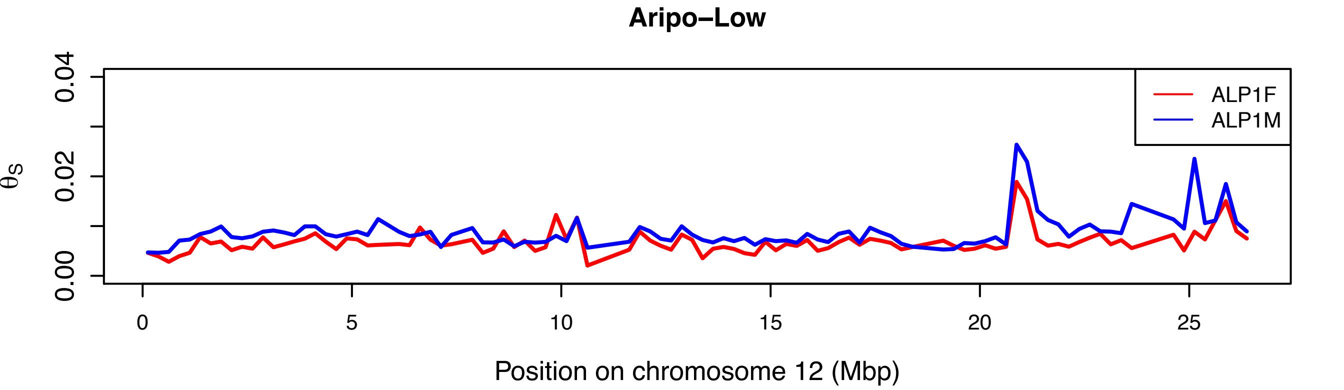
Aripo–Low
ALP1F
ALP1M
$\theta_S$
0.00
0.02
0.04
0
5
10
15
20
25
Position on chromosome 12 (Mbp)

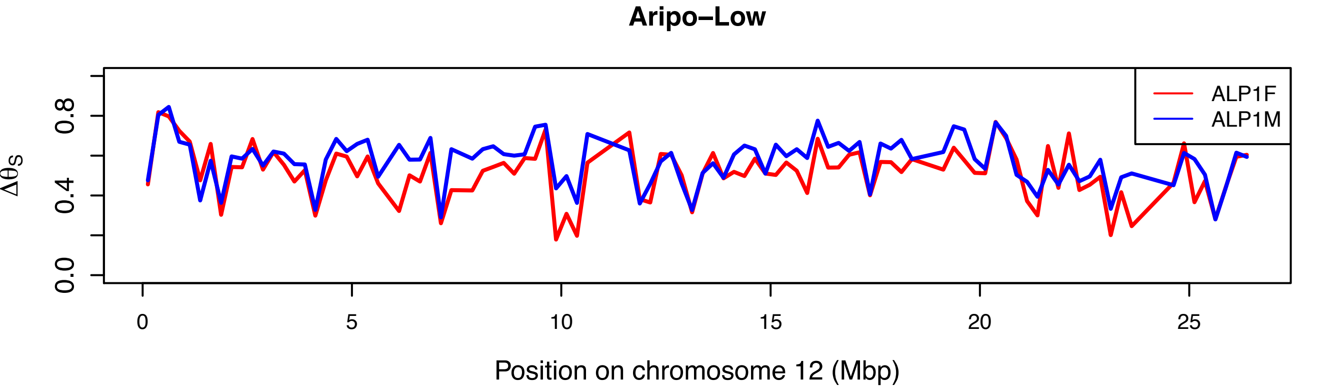
Aripo–Low
ALP1F
ALP1M
$\Delta\theta_S$
0.0
0.4
0.8
0
5
10
15
20
25
Position on chromosome 12 (Mbp)

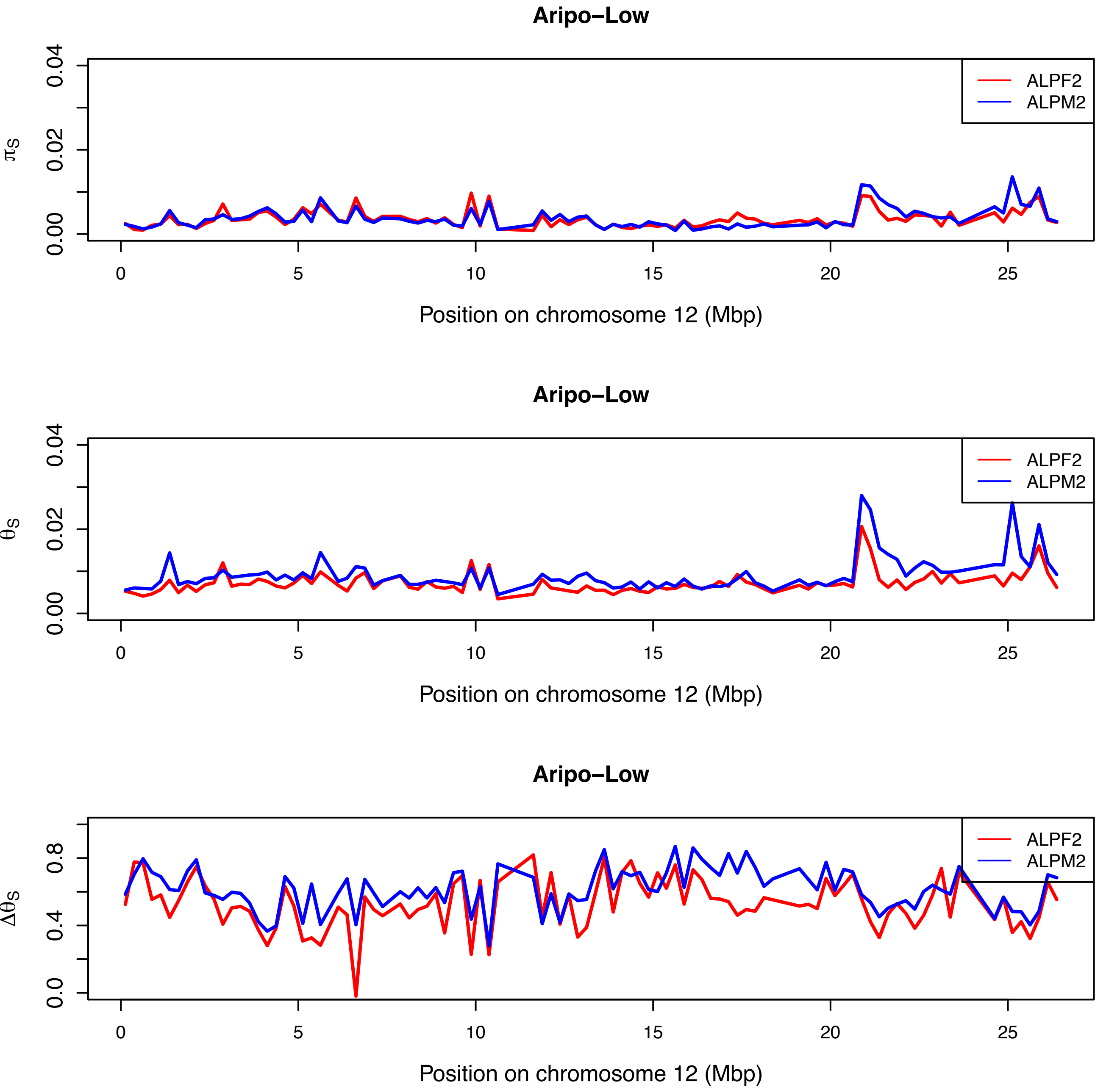

Aripo–Low
ALPF2
ALPM2
$\pi_S$
0.00
0.02
0.04
0
5
10
15
20
25
Position on chromosome 12 (Mbp)
Aripo–Low
ALPF2
ALPM2
$\theta_S$
0.00
0.02
0.04
0
5
10
15
20
25
Position on chromosome 12 (Mbp)
Aripo–Low
ALPF2
ALPM2
$\Delta\theta_S$
0.0
0.4
0.8
0
5
10
15
20
25
Position on chromosome 12 (Mbp)

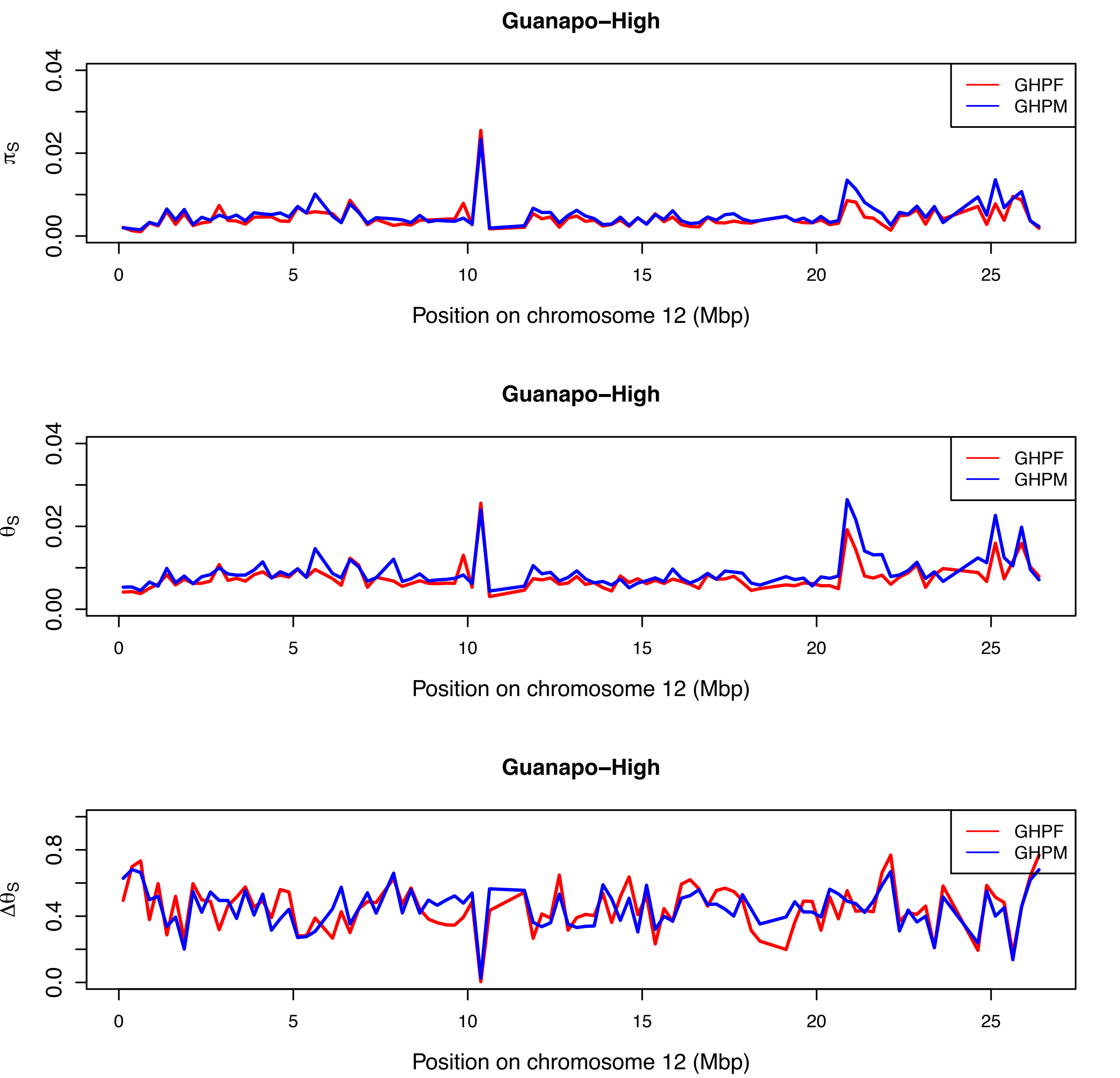

Guanapo–High
$\pi_S$
0.00
0.02
0.04
GHPF
GHPM
0
5
10
15
20
25
Position on chromosome 12 (Mbp)
Guanapo–High
$\theta_S$
0.00
0.02
0.04
GHPF
GHPM
0
5
10
15
20
25
Position on chromosome 12 (Mbp)
Guanapo–High
$\Delta\theta_S$
0.0
0.4
0.8
GHPF
GHPM
0
5
10
15
20
25
Position on chromosome 12 (Mbp)

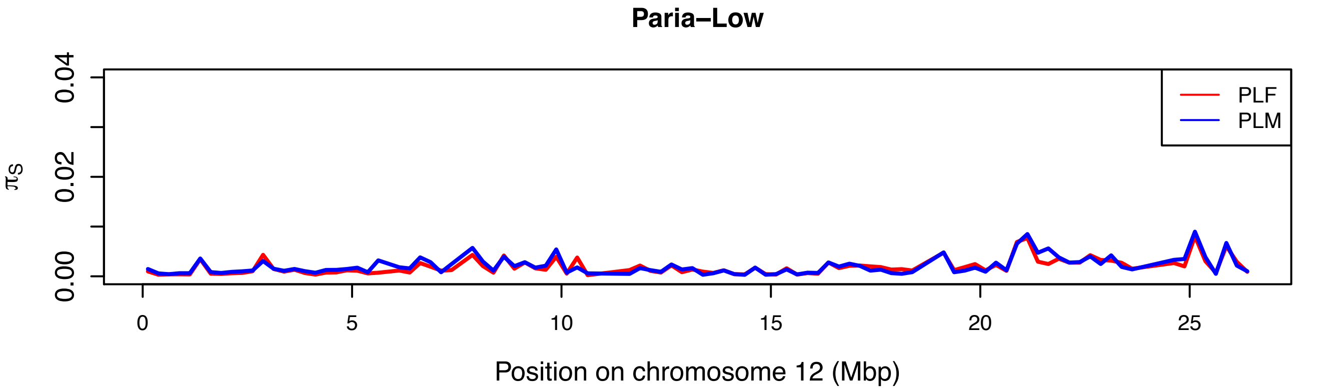
Paria–Low
$\pi_S$
0.00
0.02
0.04
PLF
PLM
0
5
10
15
20
25
Position on chromosome 12 (Mbp)

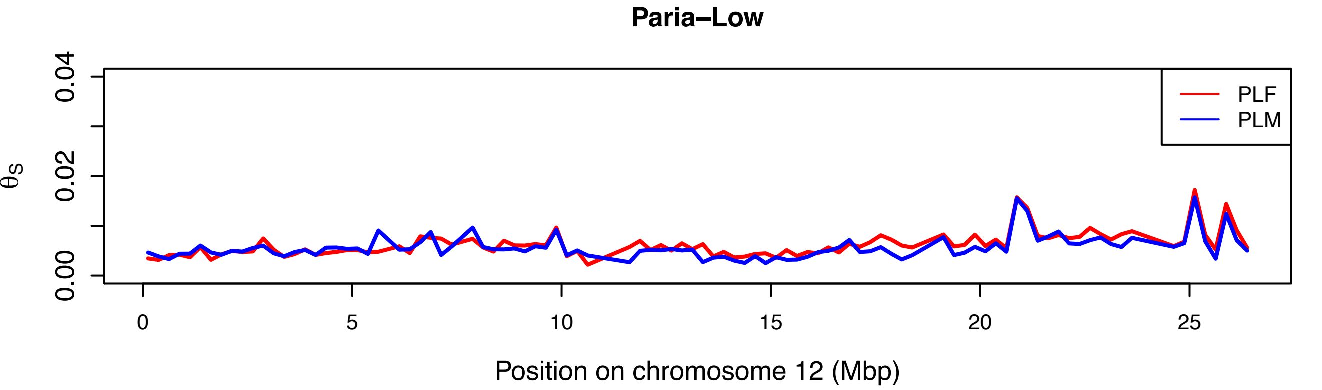
Paria–Low
$\theta_S$
0.00
0.02
0.04
PLF
PLM
0
5
10
15
20
25
Position on chromosome 12 (Mbp)

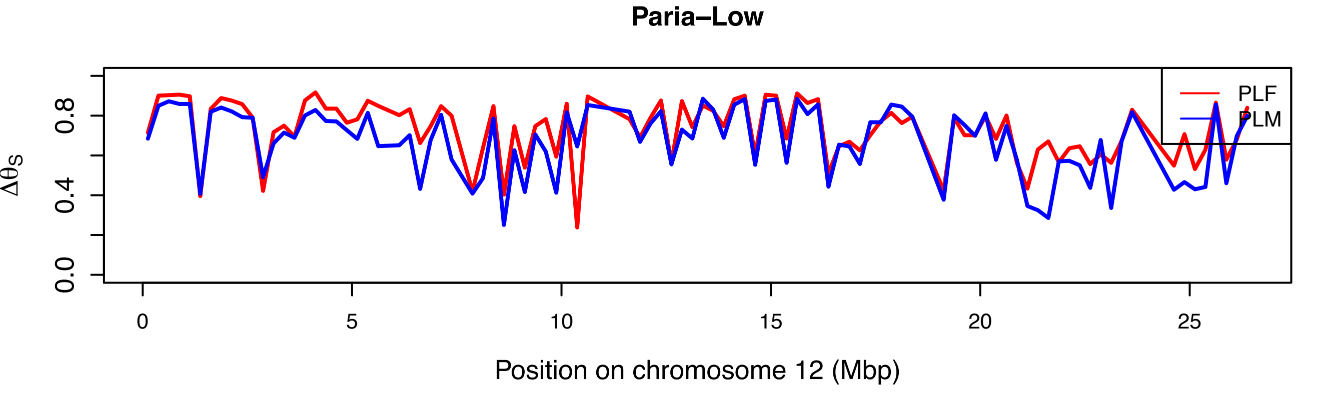
Paria–Low
$\Delta\theta_S$
0.0
0.4
0.8
PLF
PLM
0
5
10
15
20
25
Position on chromosome 12 (Mbp)

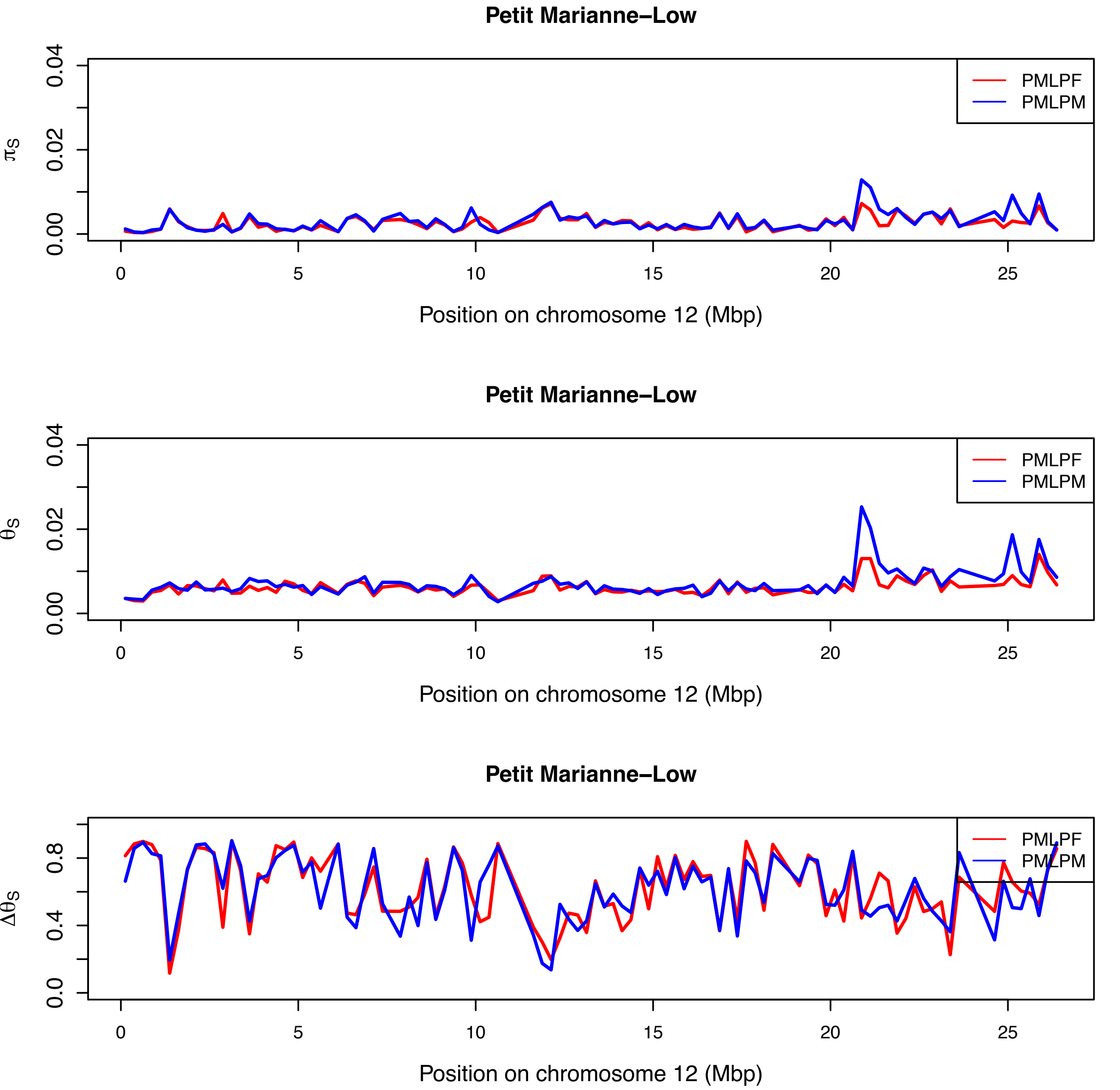

Petit Marianne–Low
$\pi_S$
0.00
0.02
0.04
PMLPF
PMLPM
0
5
10
15
20
25
Position on chromosome 12 (Mbp)
Petit Marianne–Low
$\theta_S$
0.00
0.02
0.04
PMLPF
PMLPM
0
5
10
15
20
25
Position on chromosome 12 (Mbp)
Petit Marianne–Low
$\Delta\theta_S$
0.0
0.4
0.8
PMLPF
PMLPM
0
5
10
15
20
25
Position on chromosome 12 (Mbp)

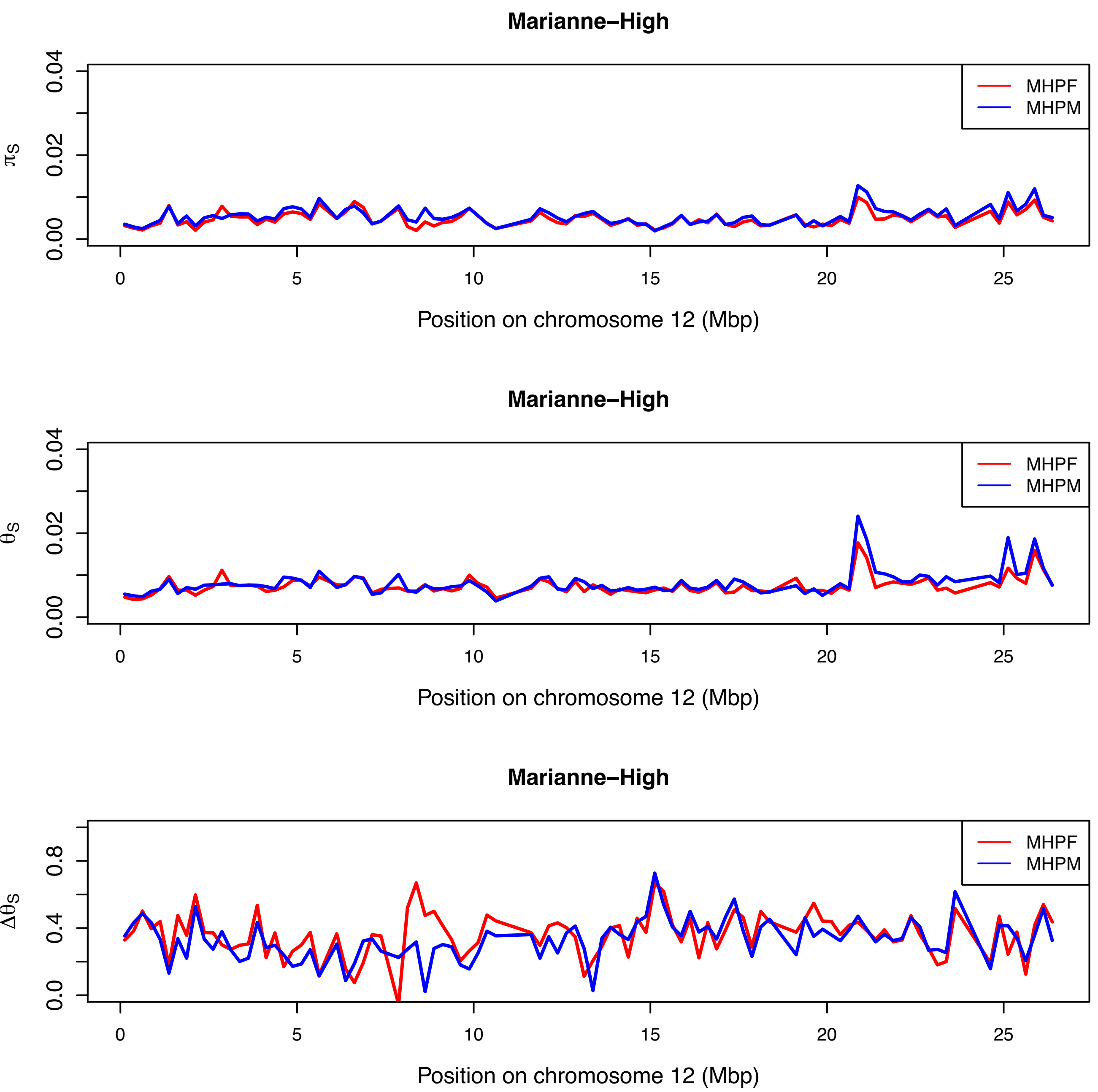
Marianne–High
0.00
0.02
0.04
πS
MHPF
MHPM
0
5
10
15
20
25
Position on chromosome 12 (Mbp)
Marianne–High
0.00
0.02
0.04
θS
MHPF
MHPM
0
5
10
15
20
25
Position on chromosome 12 (Mbp)
Marianne–High
0.0
0.4
0.8
ΔθS
MHPF
MHPM
0
5
10
15
20
25
Position on chromosome 12 (Mbp)

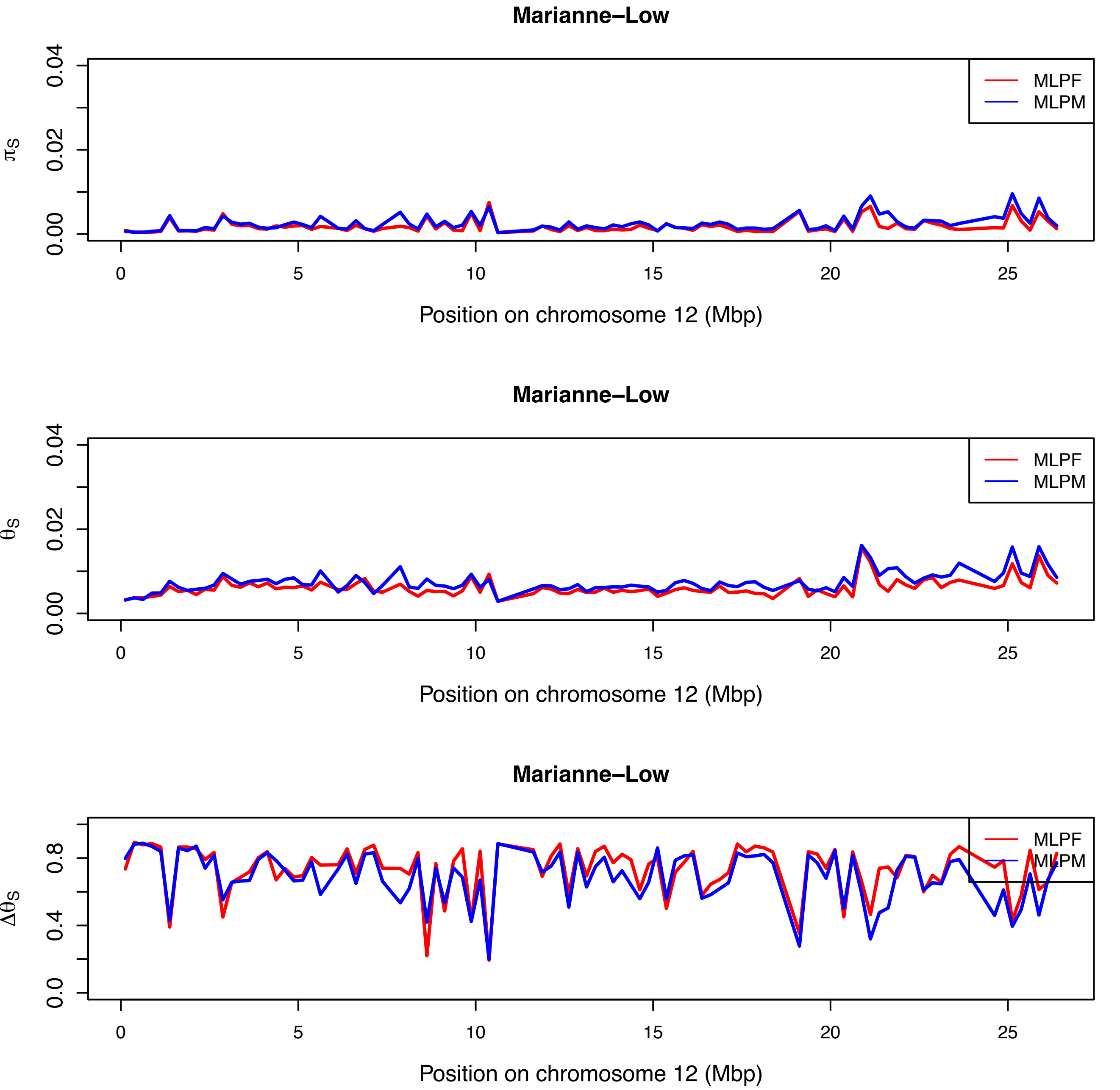
Marianne–Low
MLPF
MLPM
$\pi_S$
0.00
0.02
0.04
0
5
10
15
20
25
Position on chromosome 12 (Mbp)
Marianne–Low
MLPF
MLPM
$\theta_S$
0.00
0.02
0.04
0
5
10
15
20
25
Position on chromosome 12 (Mbp)
Marianne–Low
MLPF
MLPM
$\Delta\theta_S$
0.0
0.4
0.8
0
5
10
15
20
25
Position on chromosome 12 (Mbp)

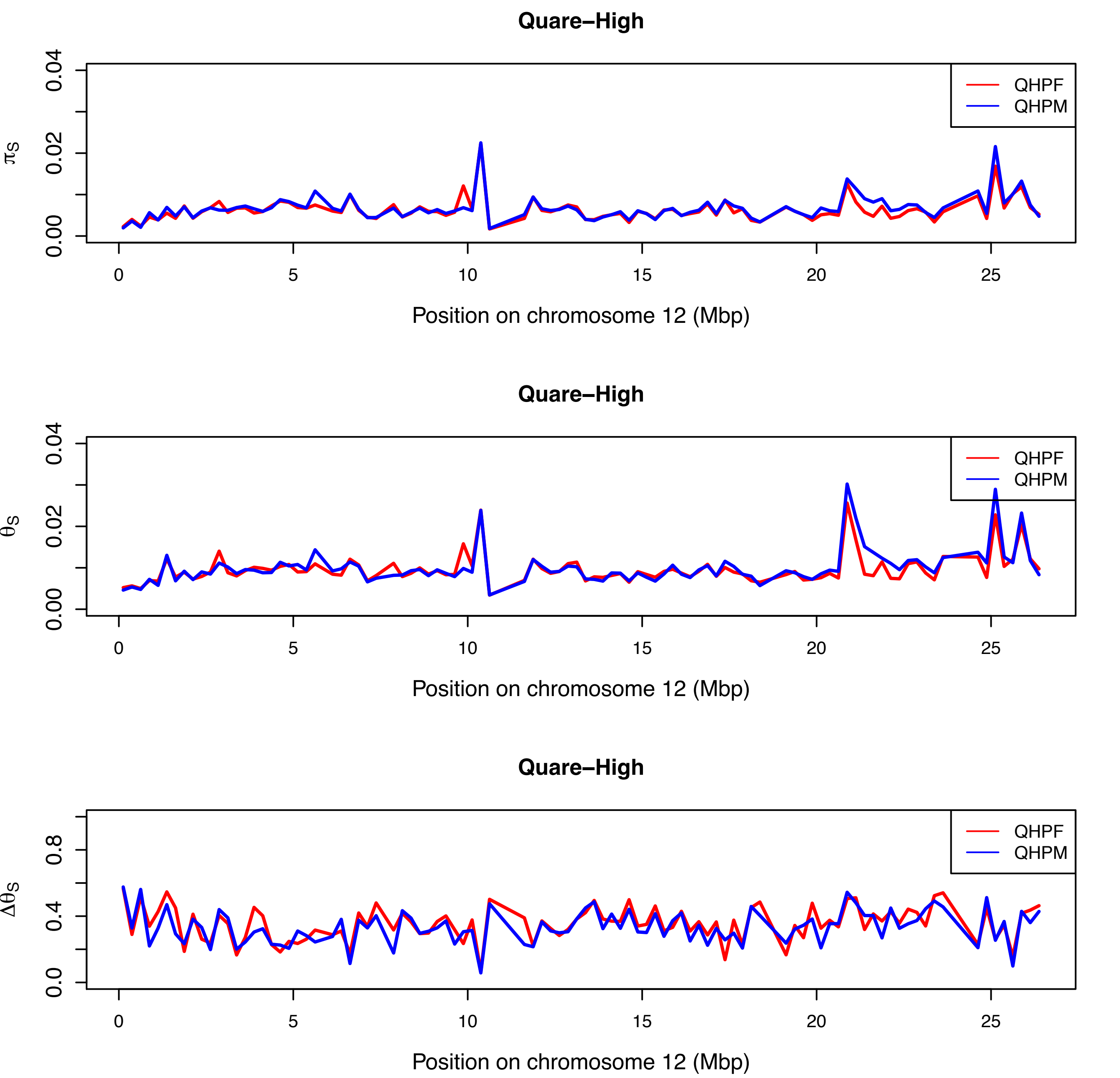
Quare–High
QHPF
QHPM
$\pi_S$
0.00
0.02
0.04
0
5
10
15
20
25
Position on chromosome 12 (Mbp)
Quare–High
QHPF
QHPM
$\theta_S$
0.00
0.02
0.04
0
5
10
15
20
25
Position on chromosome 12 (Mbp)
Quare–High
QHPF
QHPM
$\Delta\theta_S$
0.0
0.4
0.8
0
5
10
15
20
25
Position on chromosome 12 (Mbp)

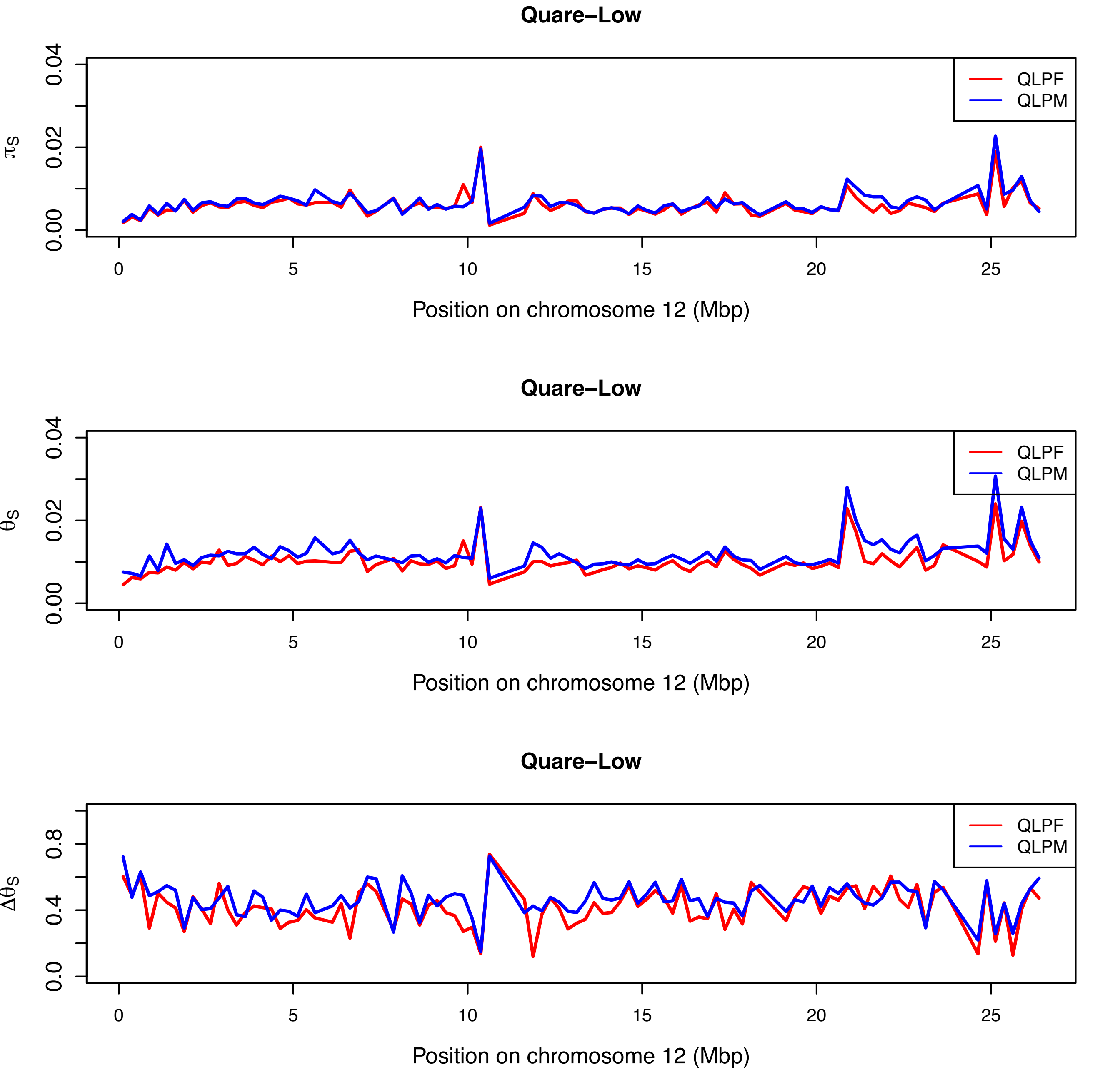
Quare−Low
$\pi_S$
0.00
0.02
0.04
QLPF
QLPM
0
5
10
15
20
25
Position on chromosome 12 (Mbp)
Quare−Low
$\theta_S$
0.00
0.02
0.04
QLPF
QLPM
0
5
10
15
20
25
Position on chromosome 12 (Mbp)
Quare−Low
$\Delta\theta_S$
0.0
0.4
0.8
QLPF
QLPM
0
5
10
15
20
25
Position on chromosome 12 (Mbp)

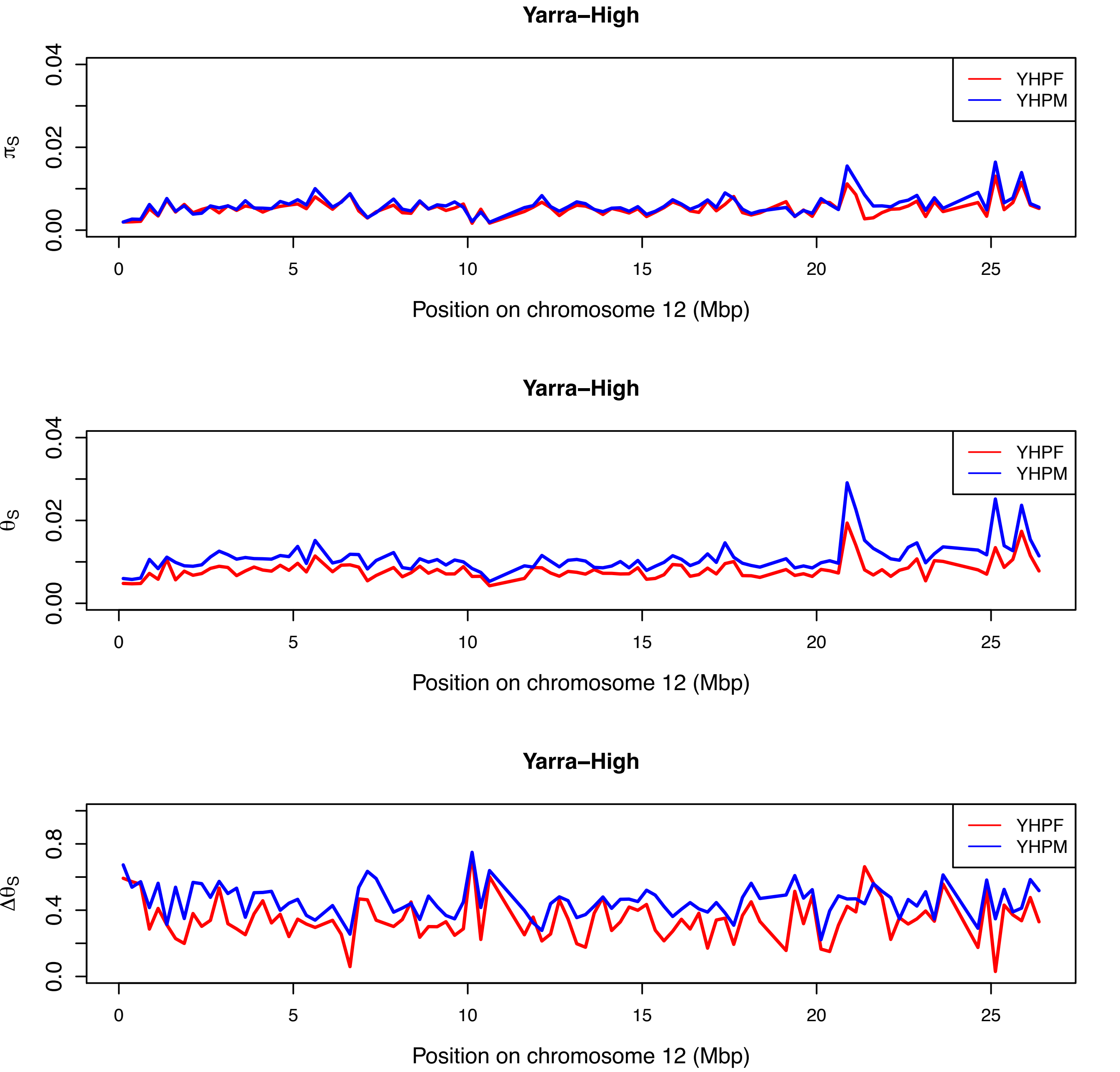

Yarra–High
YHPF
YHPM
$\pi_S$
0.00
0.02
0.04
0
5
10
15
20
25
Position on chromosome 12 (Mbp)
Yarra–High
YHPF
YHPM
$\theta_S$
0.00
0.02
0.04
0
5
10
15
20
25
Position on chromosome 12 (Mbp)
Yarra–High
YHPF
YHPM
$\Delta\theta_S$
0.0
0.4
0.8
0
5
10
15
20
25
Position on chromosome 12 (Mbp)

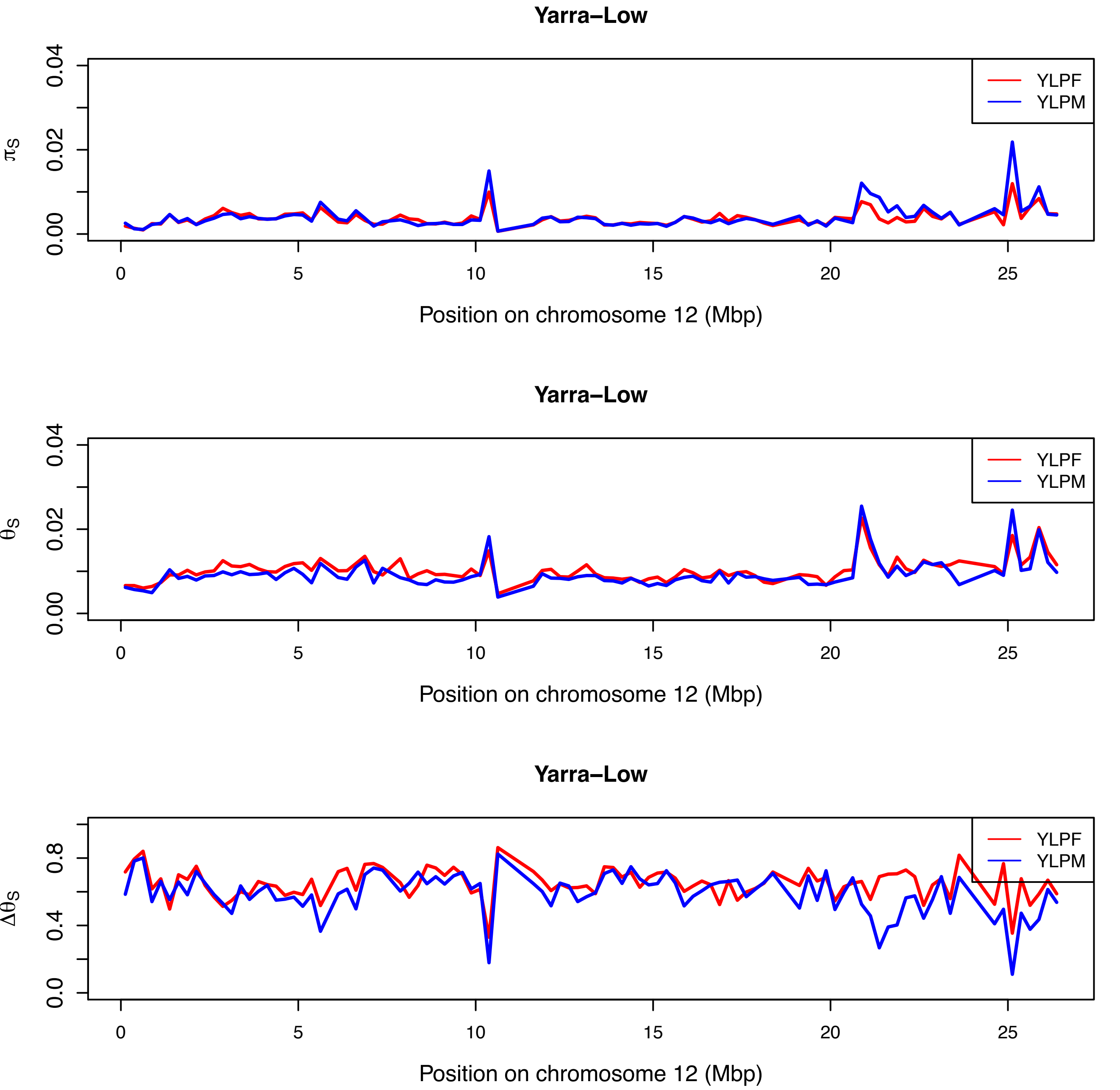

Yarra–Low
YLPF
YLPM
$\pi_S$
0.00
0.02
0.04
0
5
10
15
20
25
Position on chromosome 12 (Mbp)
Yarra–Low
YLPF
YLPM
$\theta_S$
0.00
0.02
0.04
0
5
10
15
20
25
Position on chromosome 12 (Mbp)
Yarra–Low
YLPF
YLPM
$\Delta\theta_S$
0.0
0.4
0.8
0
5
10
15
20
25
Position on chromosome 12 (Mbp)

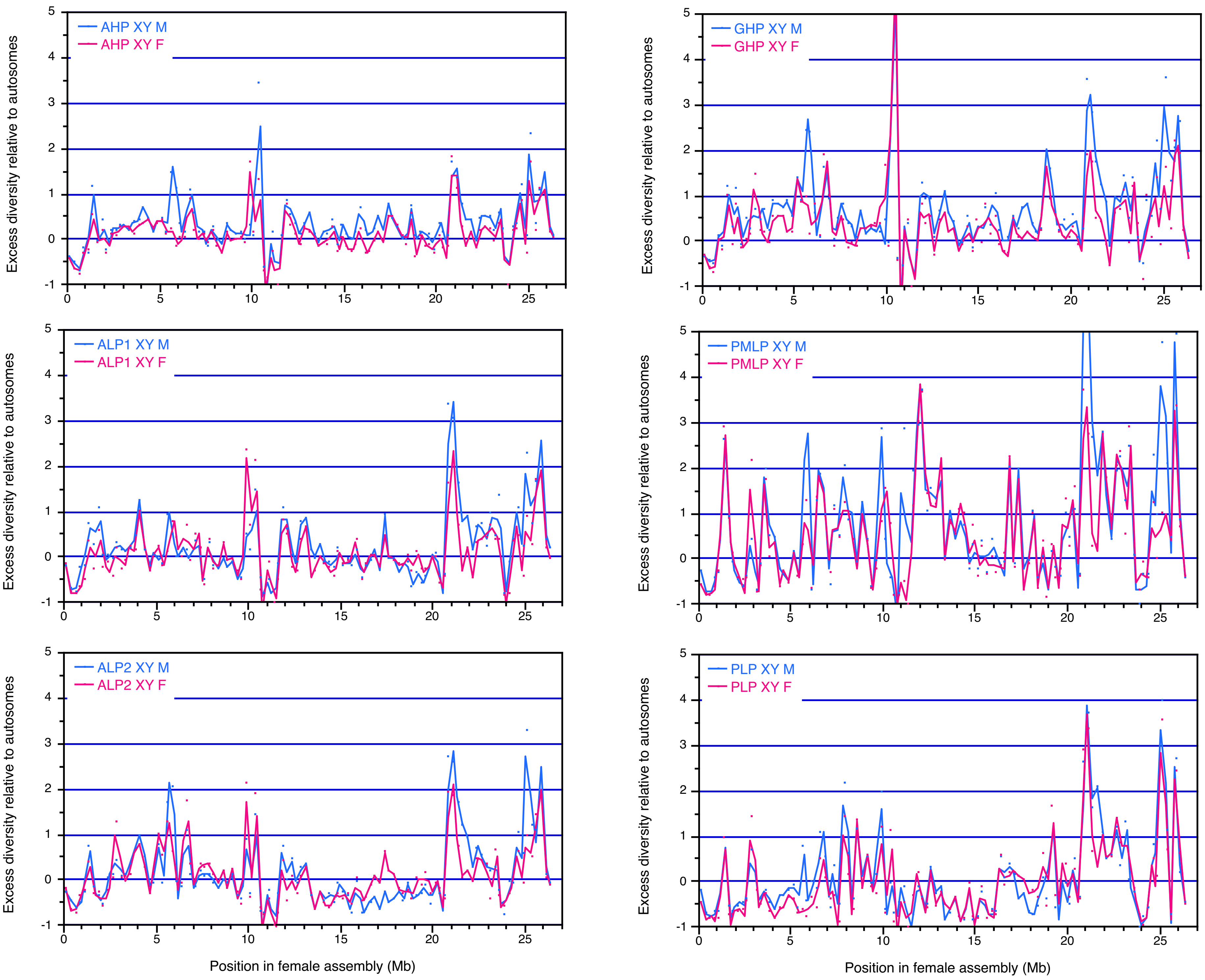

AHP XY M
AHP XY F
GHP XY M
GHP XY F
ALP1 XY M
ALP1 XY F
PMLP XY M
PMLP XY F
ALP2 XY M
ALP2 XY F
PLP XY M
PLP XY F
Excess diversity relative to autosomes
Position in female assembly (Mb)

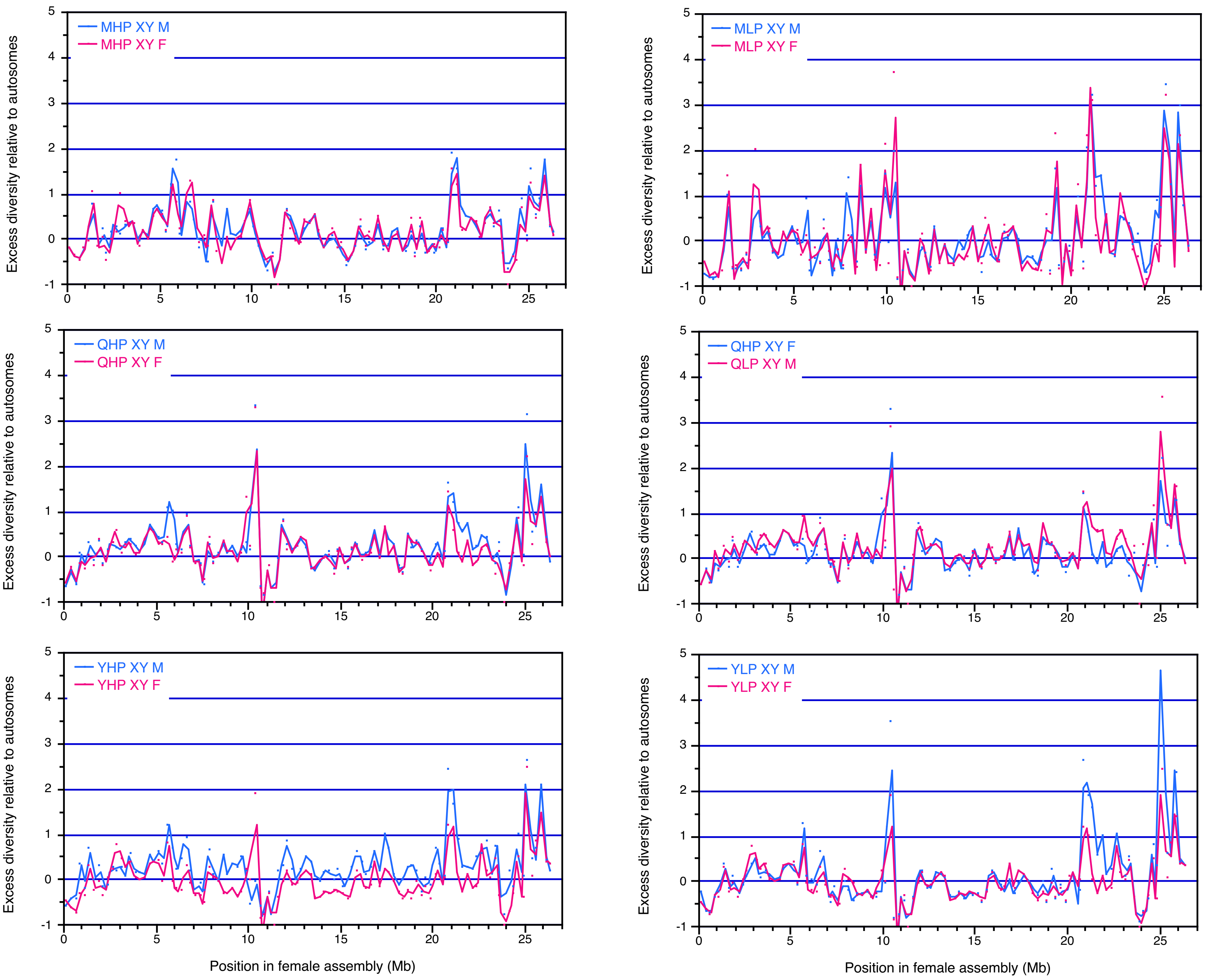

MHP XY M
MHP XY F
MLP XY M
MLP XY F
QHP XY M
QHP XY F
QHP XY F
QLP XY M
YHP XY M
YHP XY F
YLP XY M
YLP XY F
Excess diversity relative to autosomes
Position in female assembly (Mb)

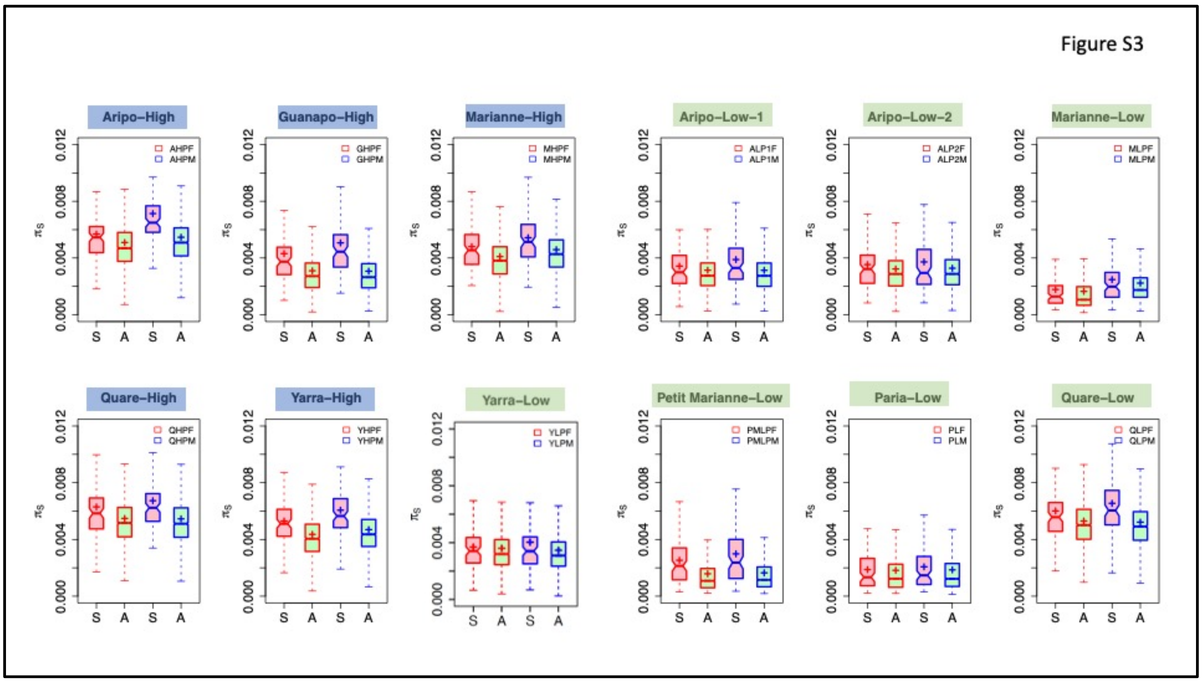


Figure S3A. Sex differences in estimated nucleotide diversity at synonymous sites (values of $\pi_S$), based on results analysed in windows (see Methods), in samples from the sites whose names are shown above the plots. High-predation (downstream) sites' names are in blue boxes, and those of low-predation, upstream sites are in green boxes. Results for females are shown in the two boxes outlined in red at the left of each plot and for males (blue outlines), on the right of each plot, and separate estimates were obtained for genes on the sex chromosome, LG12 (coloured pink boxes labelled S on the x axis) and on the autosomes (green boxes labelled A).

**Figure S4**. Associations between variants and individuals' sexes for each population sampled, for the region distal to 21 Mb of chromosome 12. The results show sites with no variation in the female samples (XFREQ= 1, labelled XP=1 in the titles of each plot), and with a range of YFREQ values (labelled YP values in the key), from the most stringent value of 1, down to the value of 0.7 shown in Figure S5.

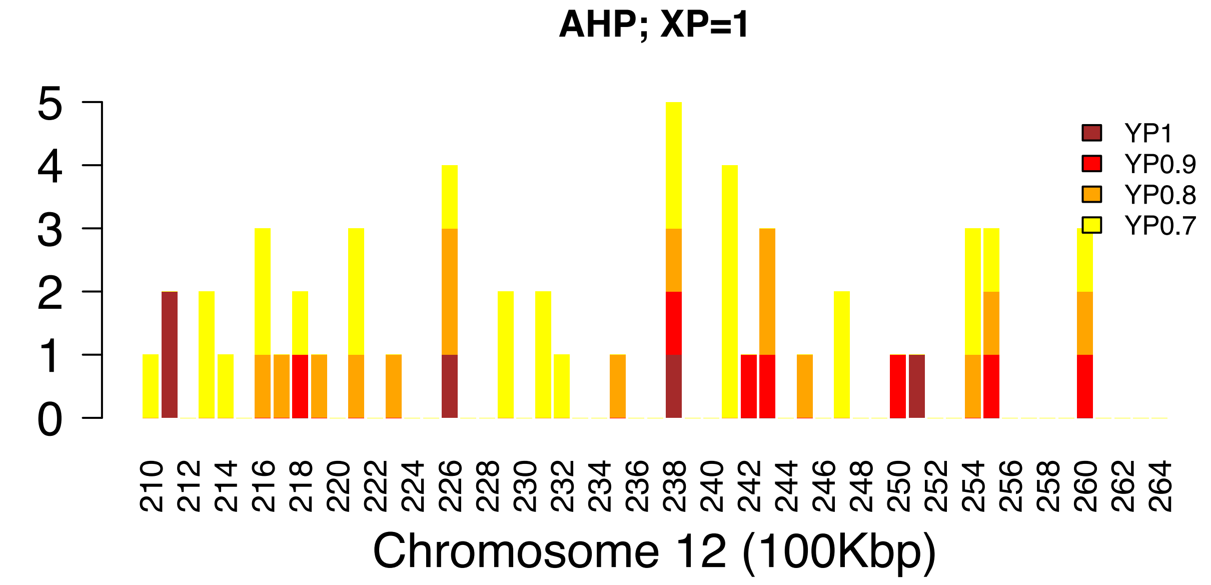
AHP; XP=1
YP1
YP0.9
YP0.8
YP0.7
Chromosome 12 (100Kbp)

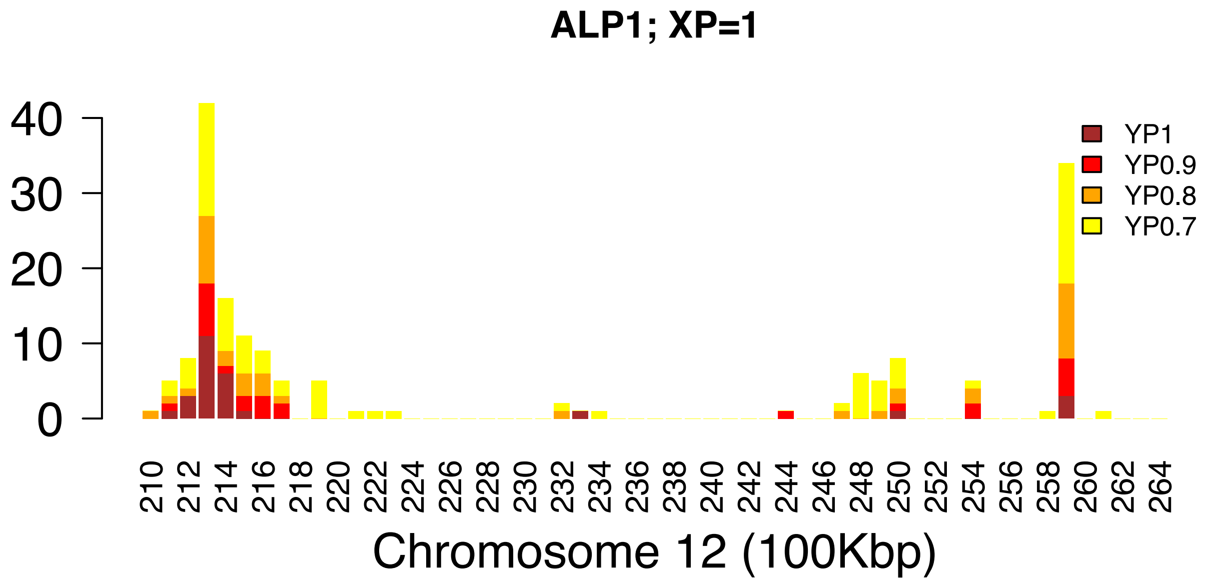
ALP1; XP=1
YP1
YP0.9
YP0.8
YP0.7
Chromosome 12 (100Kbp)

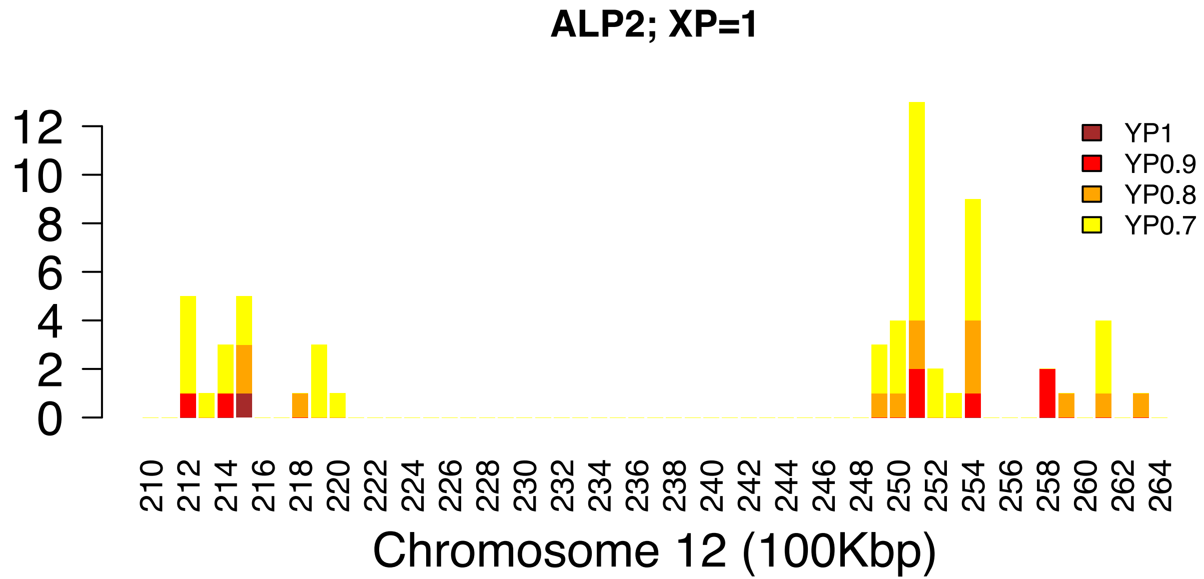
ALP2; XP=1
YP1
YP0.9
YP0.8
YP0.7
0
2
4
6
8
10
12
210 212 214 216 218 220 222 224 226 228 230 232 234 236 238 240 242 244 246 248 250 252 254 256 258 260 262 264
Chromosome 12 (100Kbp)

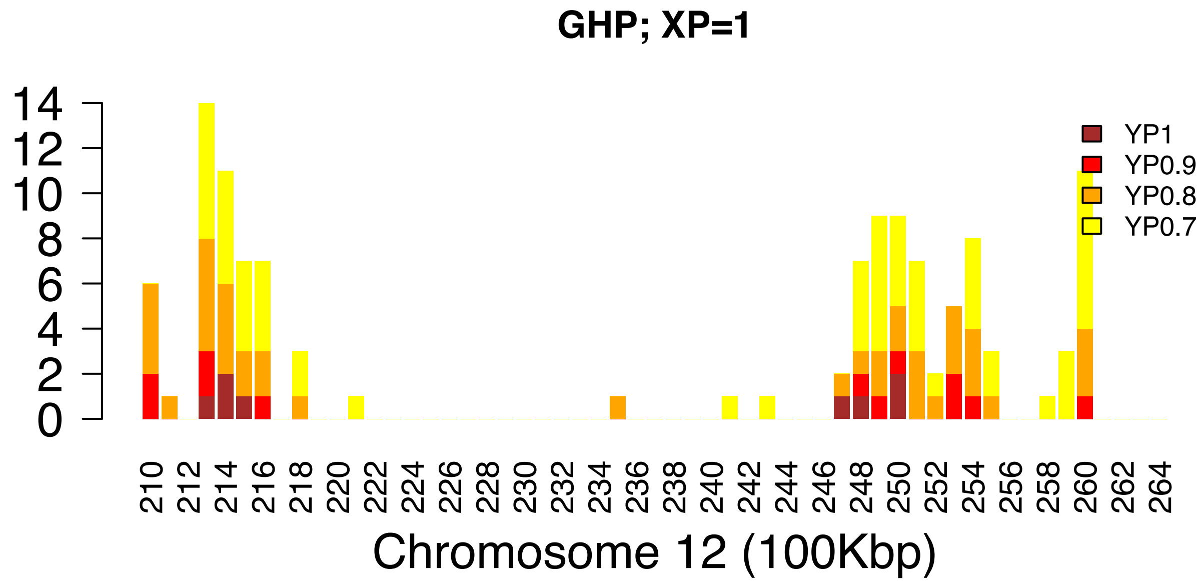
GHP; XP=1
YP1
YP0.9
YP0.8
YP0.7
0
2
4
6
8
10
12
14
210 212 214 216 218 220 222 224 226 228 230 232 234 236 238 240 242 244 246 248 250 252 254 256 258 260 262 264
Chromosome 12 (100Kbp)

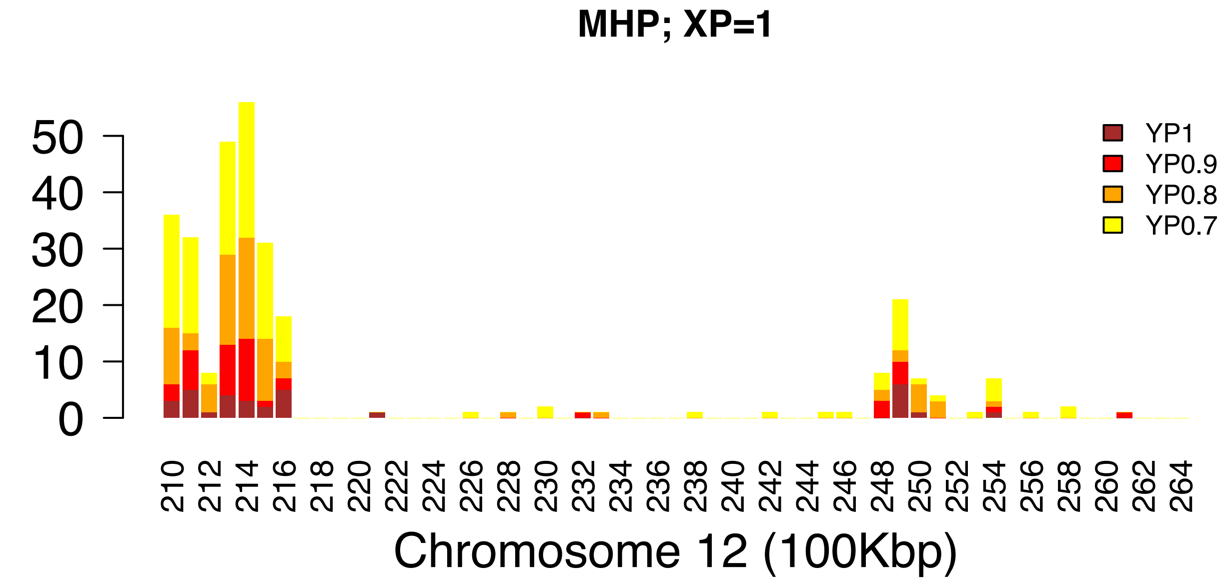
MHP; XP=1
0
10
20
30
40
50
YP1
YP0.9
YP0.8
YP0.7
210 212 214 216 218 220 222 224 226 228 230 232 234 236 238 240 242 244 246 248 250 252 254 256 258 260 262 264
Chromosome 12 (100Kbp)

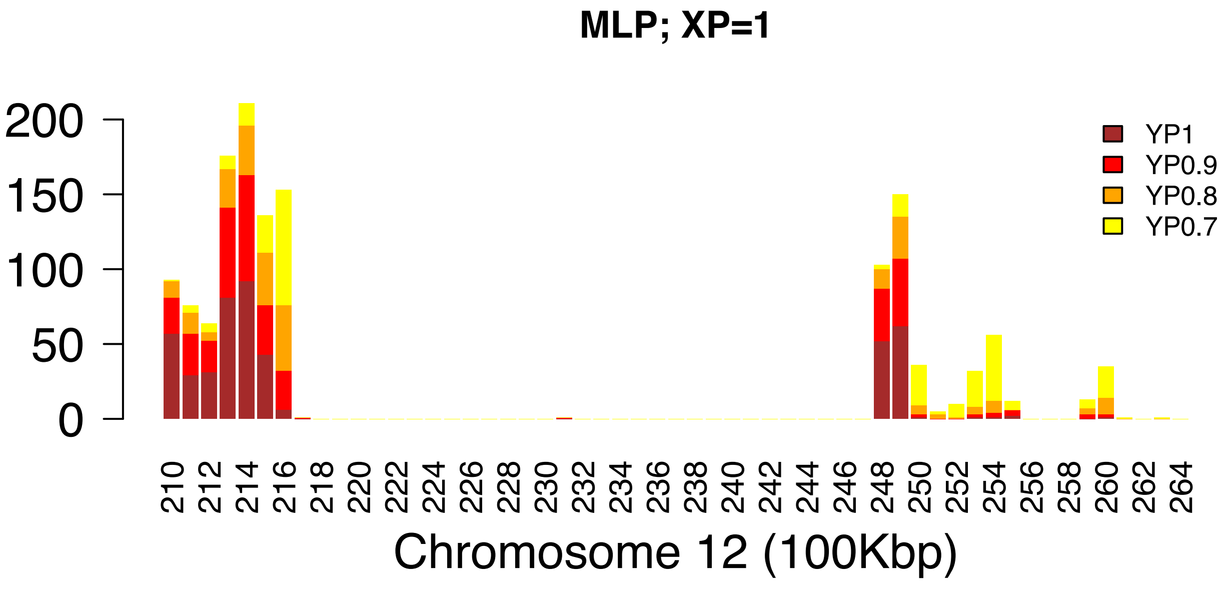
MLP; XP=1
0
50
100
150
200
YP1
YP0.9
YP0.8
YP0.7
210 212 214 216 218 220 222 224 226 228 230 232 234 236 238 240 242 244 246 248 250 252 254 256 258 260 262 264
Chromosome 12 (100Kbp)

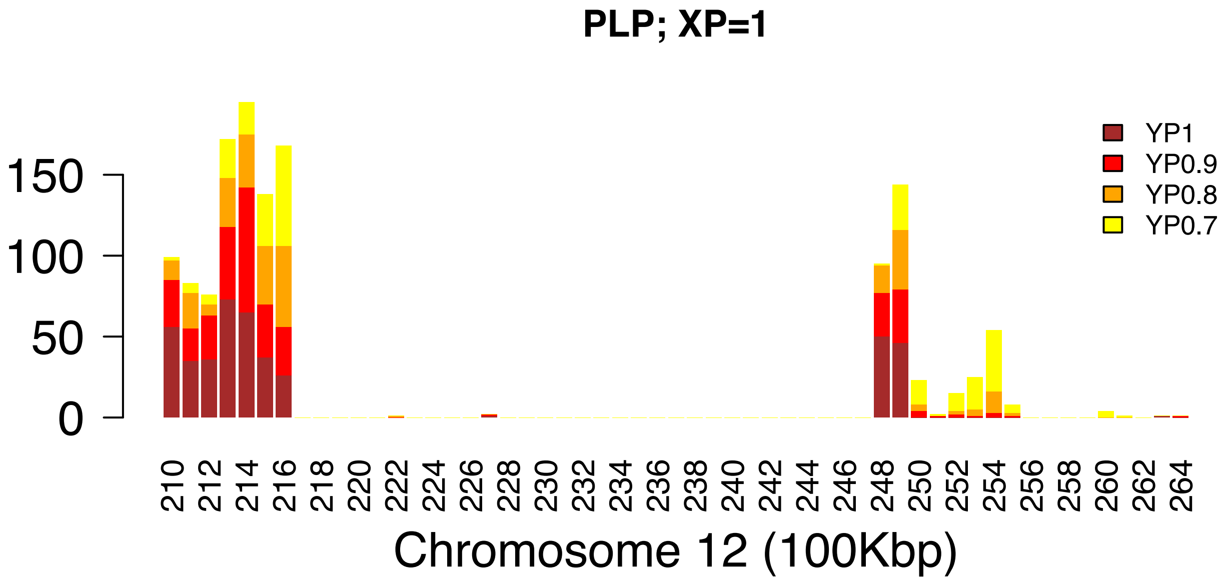
PLP; XP=1
YP1
YP0.9
YP0.8
YP0.7
Chromosome 12 (100Kbp)

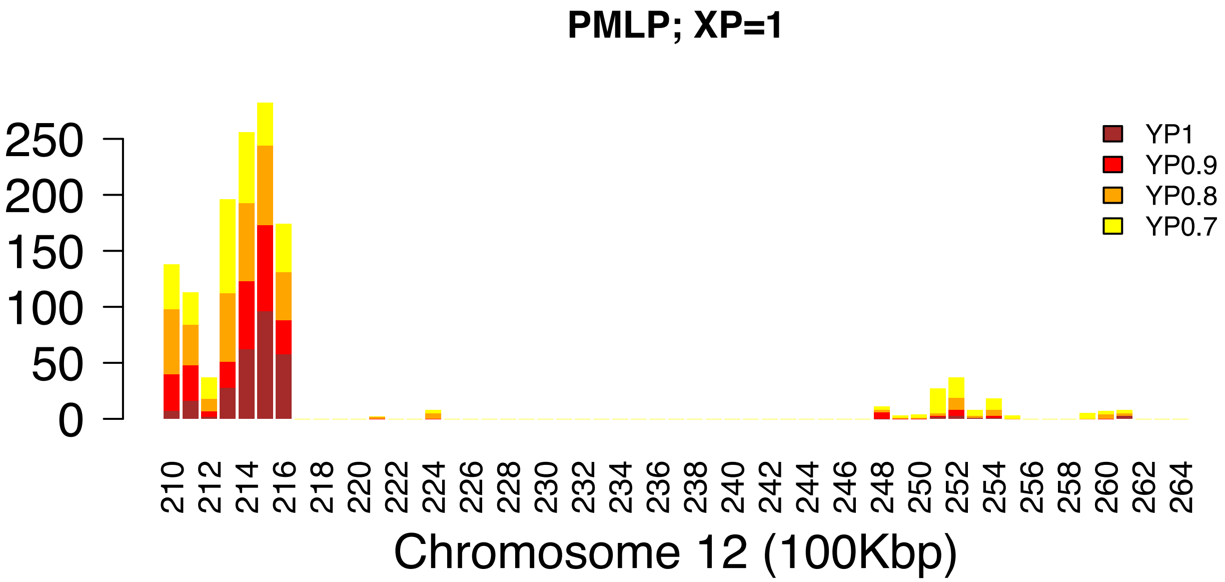
PMLP; XP=1
YP1
YP0.9
YP0.8
YP0.7
Chromosome 12 (100Kbp)

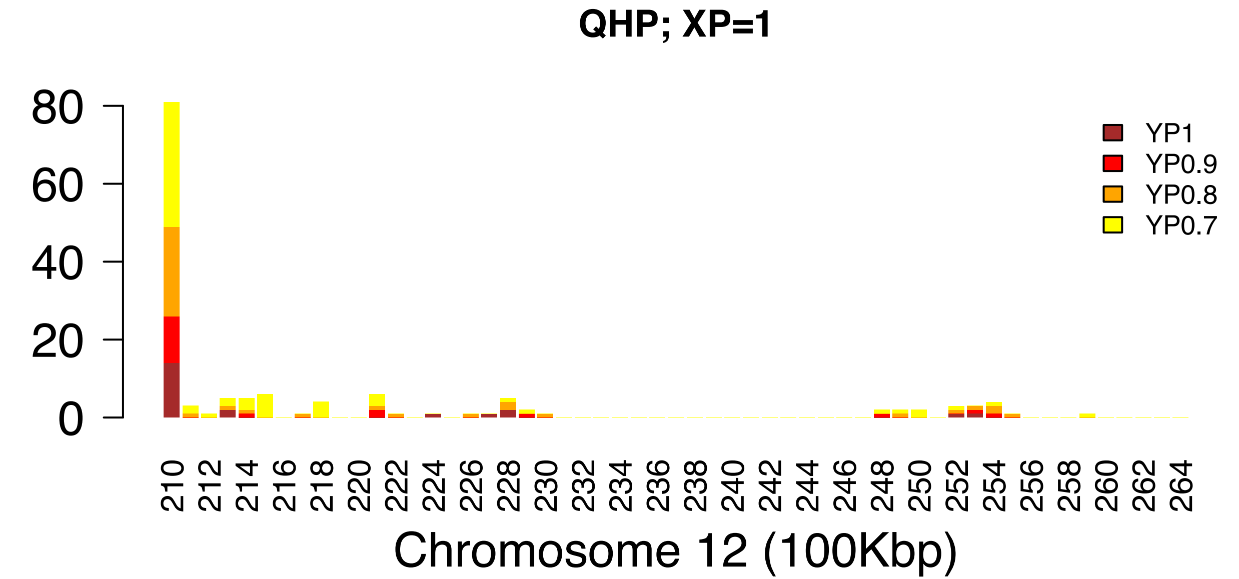
QHP; XP=1
0
20
40
60
80
YP1
YP0.9
YP0.8
YP0.7
210 212 214 216 218 220 222 224 226 228 230 232 234 236 238 240 242 244 246 248 250 252 254 256 258 260 262 264
Chromosome 12 (100Kbp)

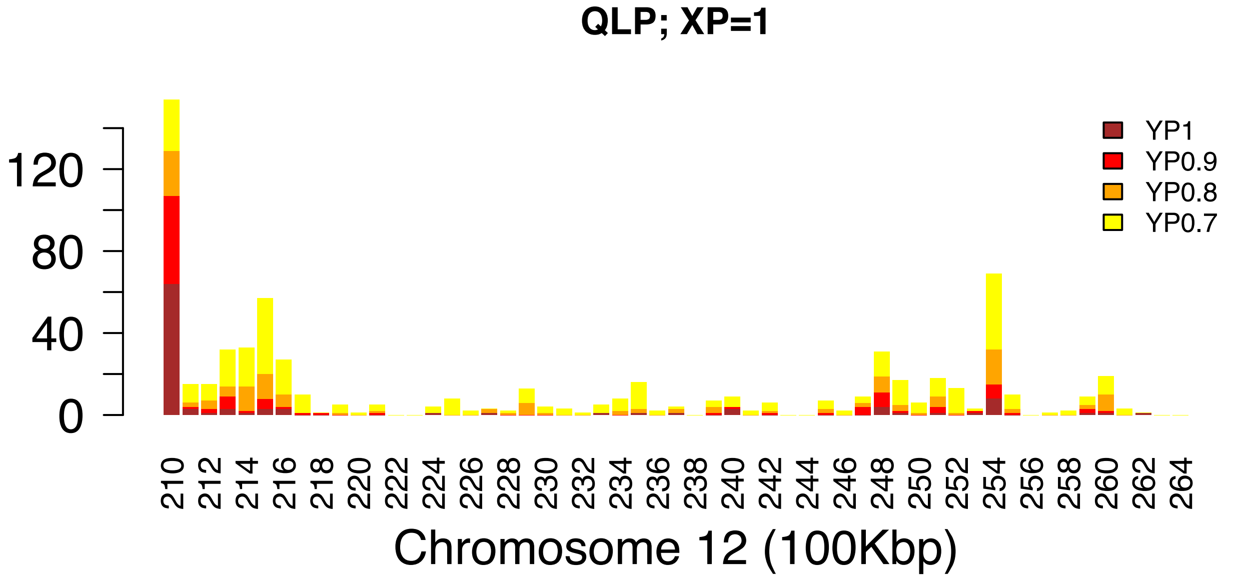
QLP; XP=1
0
40
80
120
YP1
YP0.9
YP0.8
YP0.7
210 212 214 216 218 220 222 224 226 228 230 232 234 236 238 240 242 244 246 248 250 252 254 256 258 260 262 264
Chromosome 12 (100Kbp)

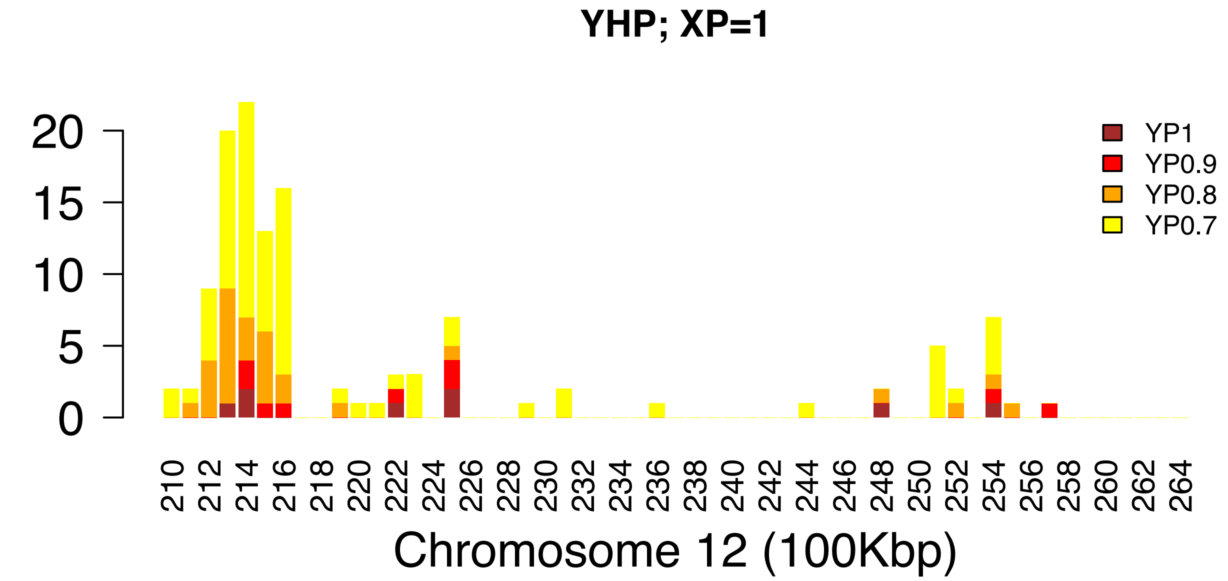
YHP; XP=1
YP1
YP0.9
YP0.8
YP0.7
0
5
10
15
20
210 212 214 216 218 220 222 224 226 228 230 232 234 236 238 240 242 244 246 248 250 252 254 256 258 260 262 264
Chromosome 12 (100Kbp)

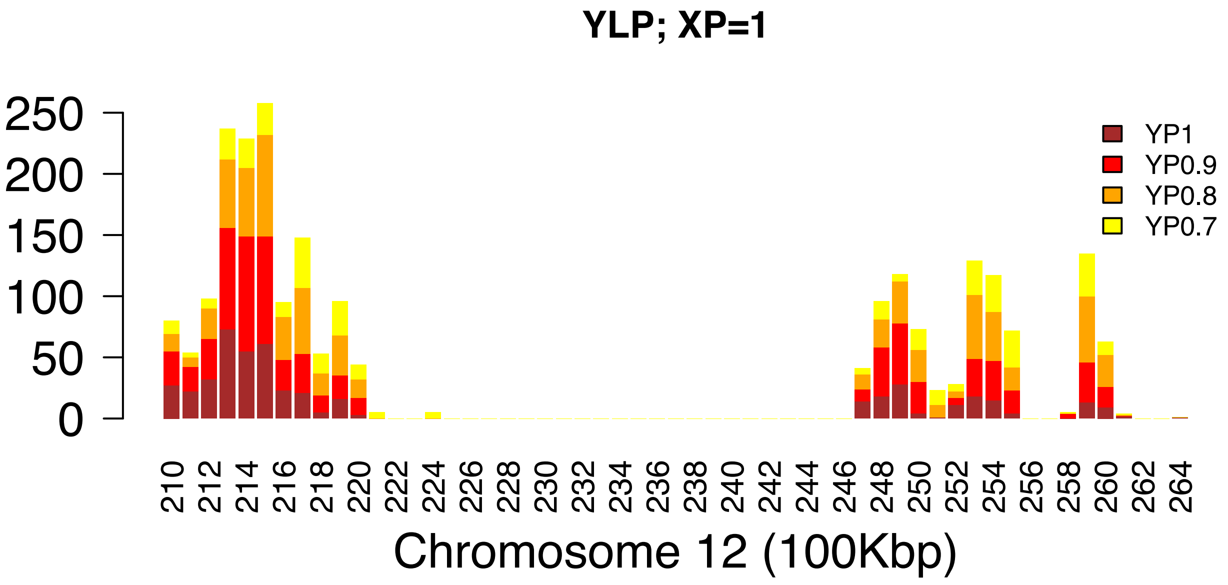
YLP; XP=1
YP1
YP0.9
YP0.8
YP0.7
0
50
100
150
200
250
210 212 214 216 218 220 222 224 226 228 230 232 234 236 238 240 242 244 246 248 250 252 254 256 258 260 262 264
Chromosome 12 (100Kbp)

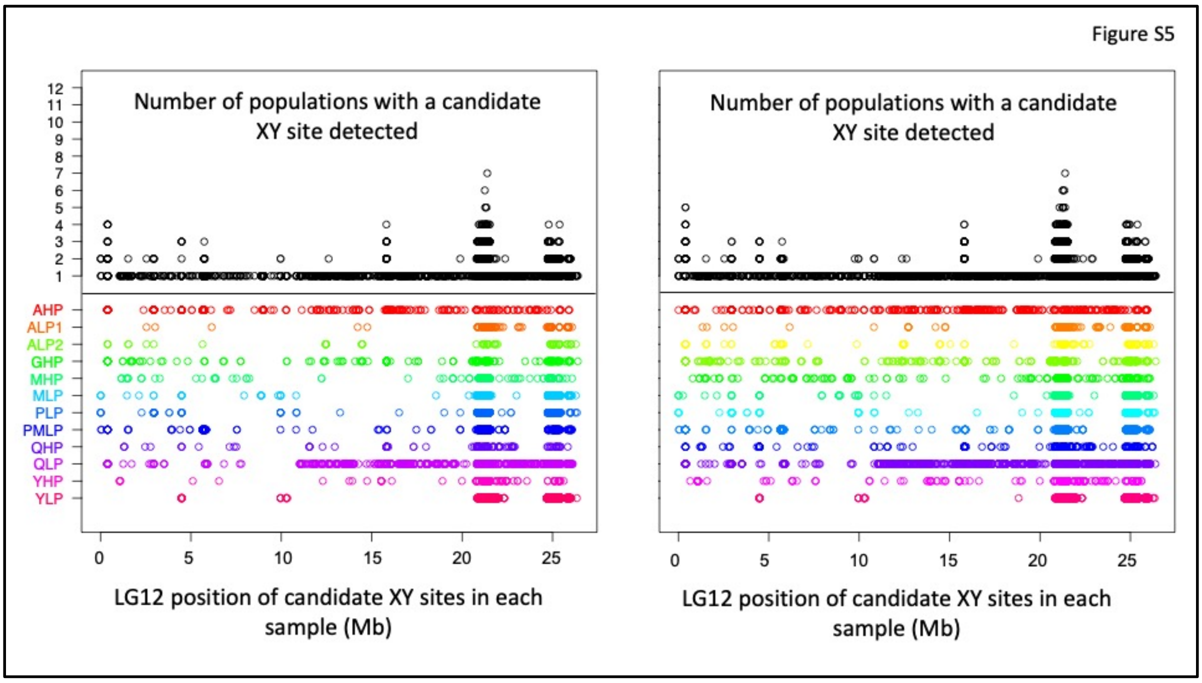


Figure S5. Candidate fully sex-linked sites in the entire LG12, analysed and displayed in the same way as in Figure 5 of the main text, but with YFREQ values > 0.7, instead of 0.9.

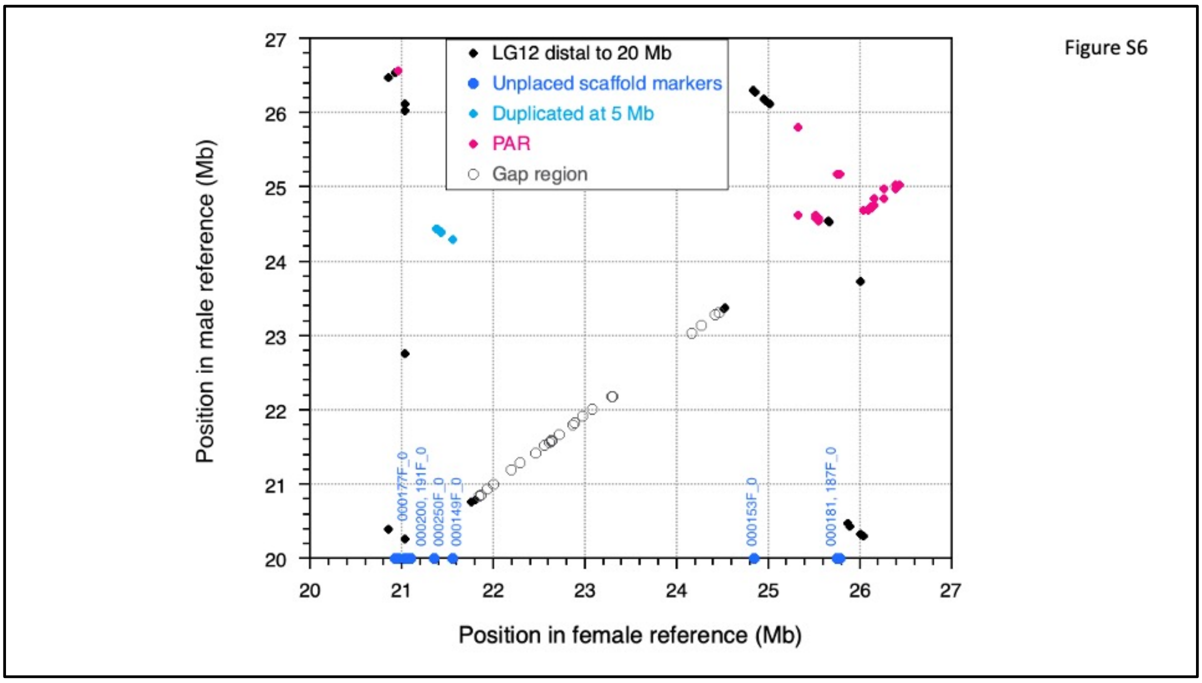


Figure S6. Comparison of positions of sites on the guppy sex chromosome in the male and female assemblies. The plot shows distal to the region that was inverted in the female assembly (see Figure S7), and it sites that have been genetically mapped to different regions in male meiosis are indicated in different colours. Markers within sequences in the gap region shown in Figures 5 and S5 are shown as open circles, and all markers mapped in this region show complete sex linkage in the sires of both families mapped. 25 markers map to the PAR in at least one of the sires studied, and 16 map to a region, contig IV, that is also duplicated at 5 Mb (Fraser et al., 2020). The plot also shows positions in the female assembly of several scaffolds that were unplaced in the male assembly, but that map genetically to this part of the guppy sex chromosomes in our sires.

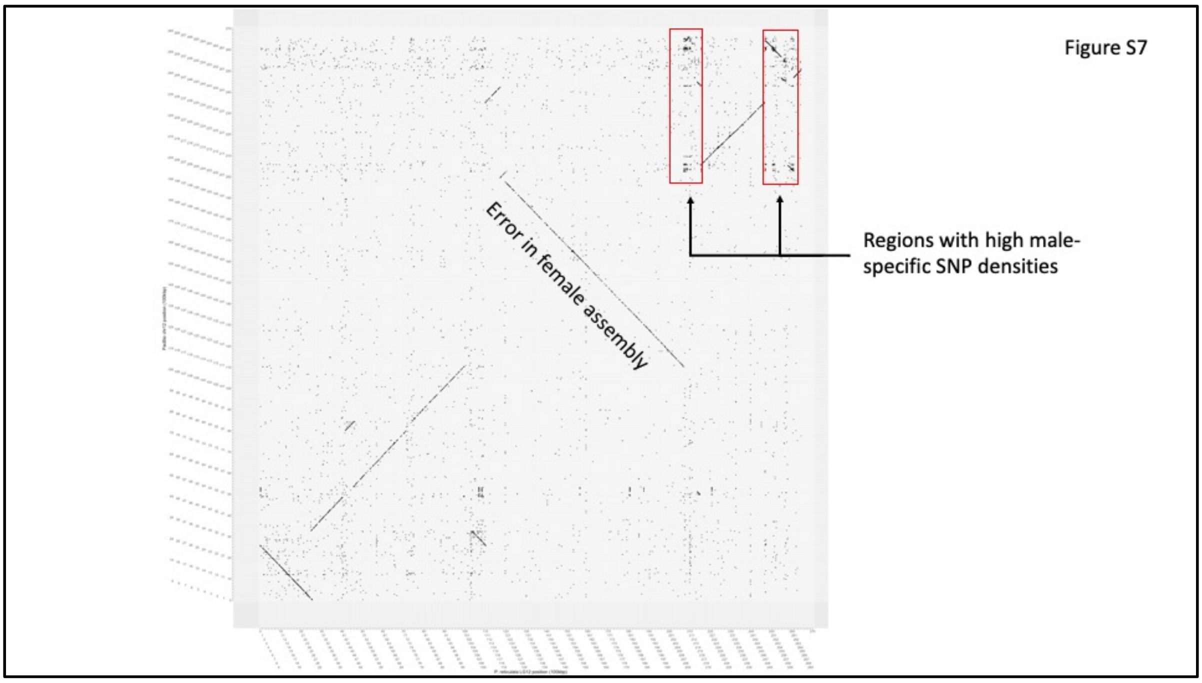


Figure S7. Comparison of the female (x axis) and male (y axis) assemblies of the guppy sex chromosome. High densities of repetitive sequences can be seen at 10 Mb, and in both regions that include most of the male-specific SNPs (se Figures 5 and S5).

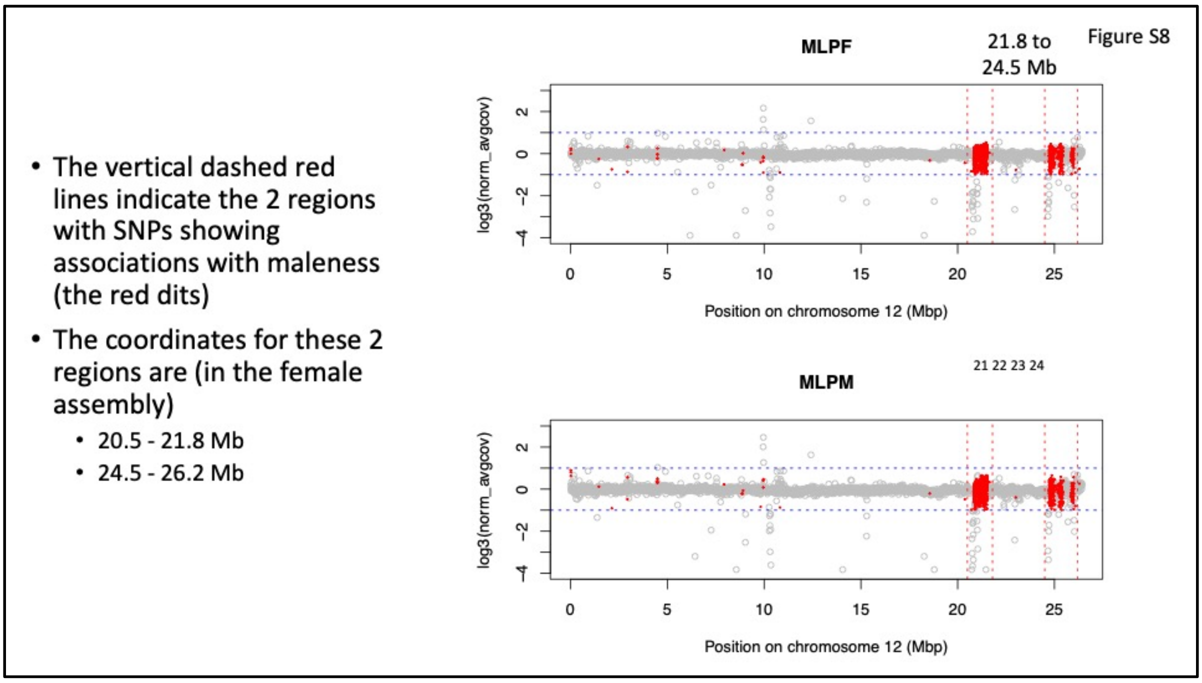


Figure S8. Mean coverage of sites in 10 kilobase windows across the sex chromosome in males and females in our pooled sequence data, with the windows where male-specific SNPs were detected shown in red (using the same threshold as for Figure S5, Yfreq=0.7), and other polymorphic sites in grey. The vertical dashed red lines indicate the two regions with high densities of SNPs showing associations with maleness; the coordinates for these regions (in the female assembly) are 20.5 - 21.8 Mb and 24.5 - 26.2 Mb. The figure shows the Marianne River low predation samples, as an example. Coverage results from other rivers are very similar.
